# GDF15 mediates inflammation-associated bone loss through a brain-bone axis

**DOI:** 10.1101/2023.11.22.568290

**Authors:** Renée Van der Cruyssen, Jan Devan, Irina Heggli, Dominik Burri, Djoere Gaublomme, Ivan Josipovic, Emilie Dumas, Carolien Vlieghe, Maria Gabriella Raimondo, Pavel Zakharov, Peggy Jacques, Sophie De Mits, Zuzanna Lukasik, Marnik Vuylsteke, Thomas Renson, Lisa Schots, Guillaume Planckaert, Flore Stappers, Tine Decruy, Julie Coudenys, Teddy Manuello, Lars Vereecke, Ruslan I. Dmitriev, Stijn Lambrecht, Luc Van Hoorebeke, Jo Lambert, Kodi Ravichandran, Andreas Ramming, Stefan Dudli, Georg Schett, Eric Gracey, Dirk Elewaut

**Affiliations:** Unit for Molecular Immunology and Inflammation, VIB-UGent Center for Inflammation Research, 9000 Ghent, Belgium; Department of Internal Medicine, Ghent University, 9000 Ghent, Belgium; Center of Experimental Rheumatology, Department of Rheumatology, University Hospital Zurich, University of Zurich, 8091 Zurich, Switzerland; Department of Physical Medicine and Rheumatology, Balgrist University Hospital, Balgrist Campus, University of Zurich, 8008 Zurich, Switzerland; Computational and Systems Biology, Biozentrum University of Basel, CH-4056 Basel, Switzerland; Swiss Institute of Bioinformatics, CH-4056 Basel, Switzerland; UGCT-Department of Physics and Astronomy, Ghent University, 9000 Ghent, Belgium; Department of Internal Medicine 3—Rheumatology and Immunology, Friedrich-Alexander-Universität Erlangen-Nürnberg (FAU) and Universitätsklinikum, 91054 Erlangen, Germany; Deutsches Zentrum Immuntherapie, Friedrich-Alexander-Universität Erlangen-Nürnberg (FAU) and Universitätsklinikum, 91054 Erlangen, Germany10^9^ Division of Immunobiology, Department of Pathology and; Immunology, Washington University School of Medicine, St. Louis, MO 63110, USA; Department of Rheumatology, Ghent University Hospital, 9000 Ghent, Belgium; Gnomixx, 9090 Melle, Belgium; Department of Dermatology, Ghent University, 9000 Ghent, Belgium; Department of Orthopedics and Traumatology, Ghent University hospital, 9000 Ghent, Belgium; Tissue Engineering and Biomaterials Group, Department of Human Structure and Repair, Faculty of Medicine and Health Sciences, Ghent University, 9000 Ghent, Belgium; Laboratory of Clinical Chemistry and Hematology, Ghent University Hospital, 9000 Ghent, Belgium; Unit for Cell Clearance in Health and Disease, VIB Center for Inflammation Research, Ghent, Belgium

**Keywords:** GDF15, GFRAL, inflammatory arthritis, bone loss, energy metabolism, IL-23, β-adrenergic signaling, marrow adipogenic lineage precursors (MALP)

## Abstract

Metabolic mediators play an important role in regulating chronic inflammation in the body. Here we report an unexpected role for GDF15 (Growth Differentiation Factor 15), a central mediator of food intake, in inflammation-associated bone loss. GDF15 serum levels were found to be elevated in arthritis patients and inversely correlated with bone density. Despite being associated with inflammation, we found that GDF15 itself does not cause, nor contribute to, clinical or histopathological arthritis. Rather, under inflammatory conditions, GDF15 mediates trabecular bone loss through its receptor GFRAL, which is exclusively expressed in the hindbrain. GDF15-GFRAL binding results in β-adrenergic activation of MALPs (Marrow Adipocytic Lineage Precursors) in the bone marrow, which stimulate osteoclasts and trigger bone loss. These data suggest a metabolic mediator-controlled brain-bone axis in inflammation, through which bone loss is induced in a contextual rather than general manner. These findings may lead to more specific therapeutic interventions to protect bone.

## INTRODUCTION

Adipose tissue influences the function of the immune system. Obesity instigates and sustains low-grade inflammation, amplifies immune-mediated disorders and their associated comorbidities^1^. As such, factors regulating adipose tissue may aggravate inflammation and predispose patients to cardiovascular disease and metabolic comorbidities^2^.

Rheumatoid arthritis (RA) and spondyloarthritis (SpA) represent two prototypic immune-mediated inflammatory diseases that are often linked with obesity^3^. These disorders are defined by chronic inflammation of the joints leading to disability if insufficiently treated. Both forms of inflammatory arthritis are strongly linked with systemic bone loss^4–7^. It has long been assumed that bone loss in the context of inflammation is directly due to the role of inflammatory mediators, particularly cytokines^8^. Indeed, inflammation levels correlate with bone loss^9^. However, patients treated with cytokine-targeted therapies can lack adequate control of bone loss, despite effective control of arthritis^10,11^. This suggests that contrary to the current dogma, inflammation and bone loss may not be directly coupled in inflammatory arthritis.

GDF15 (Growth Differentiation Factor 15) is an emerging metabolism-associated soluble protein^12,13^. It is broadly expressed by tissues of the body, including the placenta, prostate, colon, and liver^14^. The only known receptor for GDF15 is GFRAL (GDNF (Glial Derived Neurotrophic Factor) – family receptor α-like), a molecule found uniquely in the hindbrain^15–18^. Through GFRAL, GDF15 acts as a key regulator of body weight by mediating food intake, causing weight loss. As such, GDF15 serum levels are increased in patients with obesity or metabolic syndrome, likely representing compensatory mechanisms of the body trying to limit energy uptake^19^. Its serum levels were also found to be increased in infectious and inflammatory diseases^20–22^. However, whether GDF15 actively contributes to the pathogenesis of inflammatory diseases is currently unknown^23^.

Here, we reveal an unexpected role for GDF15 in inflammatory arthritis. We observed a marked elevation of serum GDF15 in arthritis patients and its levels were found to negatively correlate with bone mineral density. We provide evidence that under steady-state conditions, elevated systemic GDF15 induces bone loss with no clinical or molecular signs of inflammation. In experimental arthritis, GDF15 plays no role in joint or extra-articular inflammation yet mediates trabecular bone loss. We thereby identified a novel brain-bone axis, by which GDF15-GFRAL triggers β-adrenergic activation of MALPs (Marrow Adipogenic Lineage Precursors) in the bone, resulting in their production of the osteoclastogenic factors, RANKL and M-CSF. These findings challenge the current dogma that bone loss in inflammatory disease results from inflammatory mediators and opens new pathways for protecting bone in the context of inflammation.

## RESULTS

### Increased GDF15 levels in arthritis associates with low bone density

We first measured circulating levels of GDF15 in human inflammatory arthritis. Examination of serum from arthritis patients showed an elevation of GDF15 in RA and SpA patients relative to healthy controls (HC) (**Figure 1A**). We found SpA patients with peripheral joint arthritis to have higher serum GDF15 levels than those with axial involvement (**Figure 1B**). In addition, SpA patients with psoriasis had higher serum GDF15 levels than those without (**Figure S1A**). In an independent cohort, we observed a similar increase in GDF15 serum levels in PsA (Psoriatic Arthritis) patients (**Figure 1C**). We also sought to determine whether GDF15 levels are linked to the degree of skin inflammation. To this end, we examined a cohort of psoriasis patients without arthritis. We observed a significant correlation between GDF15 serum levels and skin inflammation severity scores (PASI) (**Figure 1D**). Collectively, these data provide evidence that GDF15 levels are increased in various forms of immune-mediated inflammatory diseases.

**Figure 1.**
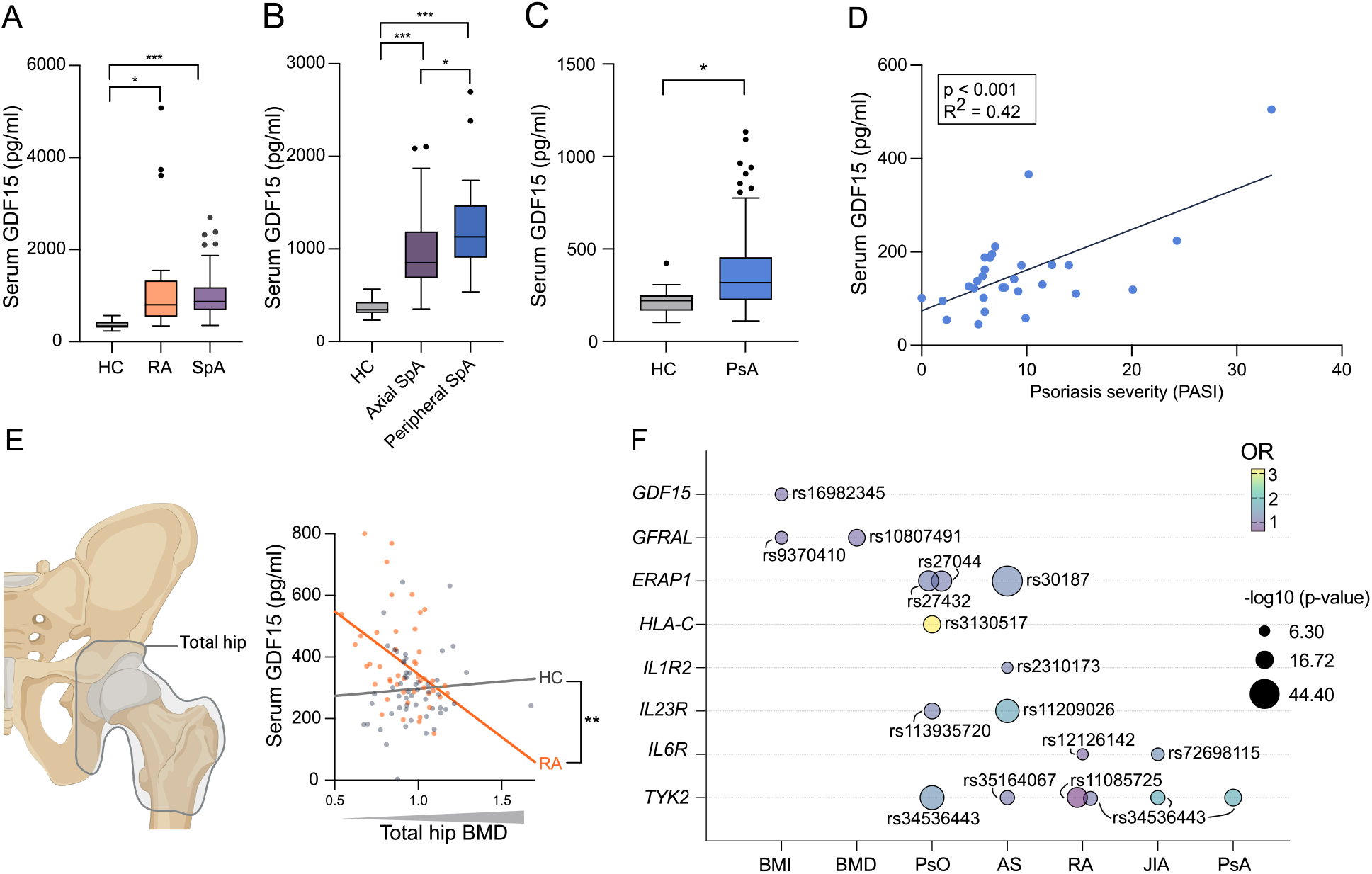
Increased GDF15 levels in arthritis associates with low bone density. (A-B) Luminex was used to measure GDF15 in serum from HC (heathy controls, n=19), RA (Rheumatoid Arthritis, n=20) and SpA (Spondyloarthritis, n=111) patients. SpA patients were subdivided based on the presence of arthritis in axial (n=71) or peripheral joints (n=17). (C) Serum GDF15 levels in an independent cohort of HC (n=31) and PsA (Psoriatic Arthritis, n=73) patients determined by ELISA. (A-C) Tukey box and whiskers plots. One-way ANOVA was used to test for differences among patient means, followed by a post-hoc Tukey test. (D) Simple linear regression was used to generate a regression line between serum GDF15 levels, determined by ELISA, and skin inflammation severity assessed using the Psoriasis Area and Severity Index (PASI) (n=29). (E) Serum GDF15 was measured by ELISA and bone mineral density (BMD) assessed by DEXA scan in RA patients (n=46) and HC (n=54). Simple linear regression with groups was used to generate regression lines between serum GDF15 and total hip BMD. The p-value refers to the significance of the difference in regression coefficients for between patient groups. (F) GWAS (Genome Wide Association Studies) disease association analysis. Each bubble shows the lead SNP (Single Nucleotide Polymorphism) in genes of interest for the implicated phenotypes or diseases. AS = Ankylosing Spondylitis, PsO = Psoriasis, JIA = Juvenile Idiopathic Arthritis. Each datapoint represents one patient, where applicable. All serum GDF15 concentrations were adjusted for age. Significances are indicated as: *p<0.05, **p<0.01, ***p<0.001.

Next, we examined whether serum GDF15 levels are associated with bone mineral density (BMD) measured by dual-energy X-ray absorptiometry (DEXA). We found a strong negative correlation between serum GDF15 levels and BMD in the hip and femoral regions rich in trabecular bone in RA patients, but not healthy controls (**Figure 1E**, **S1B, Table S1**).

We next explored publicly available genome-wide association studies (GWAS) to see if the *GDF15* or *GFRAL* loci are linked to inflammatory diseases and/or BMD. We also included BMI given GDF15’s role as a weight regulator. As controls we included genes with established pathogenic roles in inflammatory diseases, such as *TYK2*^24^. In this analysis, single nucleotide polymorphisms (SNPs) at the *GDF15/GFRAL* loci were clearly linked to BMI and/or BMD, whereas inflammatory gene SNPs were not (**Figure 1F**). Thus, the genetic link of the GDF15-GFRAL axis with bone density in the general population strongly supports a biological role for GDF15 in the regulation of bone density. We therefore focused our investigations on the *in vivo* role of GDF15 in inflammatory arthritis.

### GDF15 induces dose-dependent bone loss but not tissue inflammation

To evaluate the *in vivo* function of GDF15 we first assessed the impact of GDF15 overexpression in mice. The half-life of recombinant GDF15 in the serum is short (less than 8 hours^25^), which hampers its use in long-term studies. We circumvented this by engineering an Enhanced Episomal Vector (EEV) to express murine GDF15 under the control of a CAGs promotor (**Figure S2A**). Administering the EEV using hydrodynamic (HDD) tail vein injection results in constitutive, high levels of protein expression^26^. As a control, we used the same plasmid without the GDF15 insert (control EEV). GDF15-EEV was injected at different doses, after which the mice were closely monitored for seven weeks, with an extensive range of tissues studied at the endpoint (**Figure 2A**).

**Figure 2.**
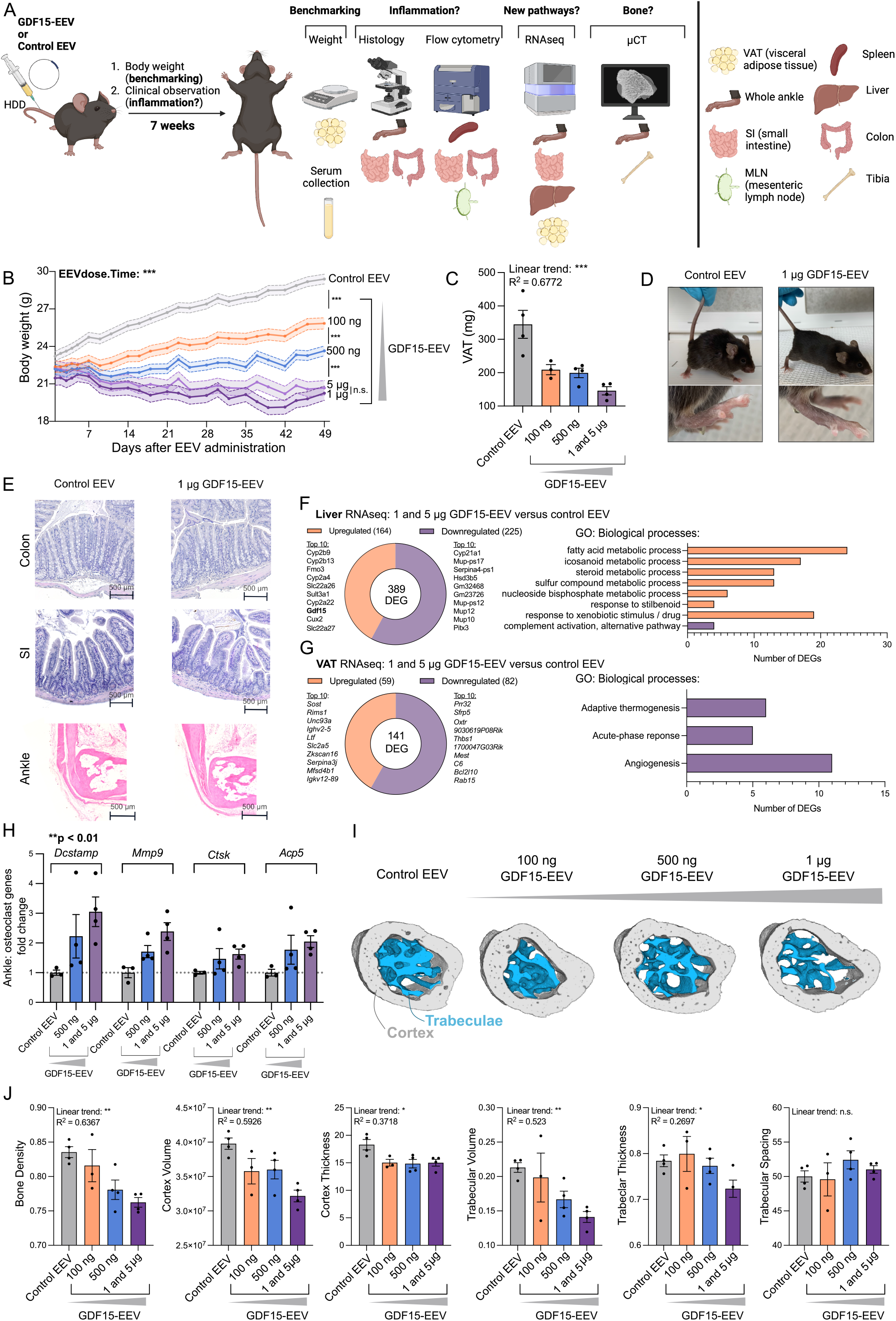
GDF15 induces dose-dependent bone loss but not tissue inflammation. (A) Schematic overview of the GDF15-Enhanced Episomal Vector (EEV) dose-response experiment. Increasing doses of GDF15-EEV or control EEV were injected using HDD (hydrodynamic) tail vein injection. The mice were weighed and clinically monitored for signs of inflammation and pain at least three times per week for seven weeks. Upon sacrifice, tissues were collected for processing as indicated in the schematic. (B) Mouse body weight over time of mice injected with control EEV or increasing doses of GDF15-EEV (n=2-4). Body weights were adjusted for differences in baseline body weight. A repeated measurements analysis was performed to assess overall changes in body weight over time between the four doses, represented by the EEVdose.Time interaction term, followed by testing pairwise contrasts between doses (control EEV versus 100ng GDF15-EEV, 100ng GDF15-EEV versus 500ng GDF15-EEV etc.). (C) Visceral adipose tissue (VAT) weight. A one-way ANOVA was used to test for differences in VAT between the doses, followed by post-hoc test for trend. (D) Mice with control EEV or 1µg GDF15-EEV display no visial signs of pain, dermatitis or arthritis. (E) Representative H&E-stained microscopy images of colon, SI (small intestine) and ankle enthesis. (F-G) Bulk RNAseq of the liver and VAT from control EEV versus 1 and 5µg GDF15-EEV treated mice. Total number of differentially expressed genes (DEGs) (log2 fold change >1.0 and <1.0, adjusted p value <0.05), and the top 10 up– and downregulated DEGs listed. Pathway analysis using GO (Gene Ontology) terms showing the biological processes (p<0.001) to which the up– or downregulated DEGs contribute. (H) Bulk RNAseq performed on ankle of GDF15-EEV treated mice. For each gene, counts normalized to control of genes displayed. A one-way ANOVA was used to test for the effect of EEV. (I) µCT images of calcaneus. Representative cross-sections shown, with the bone cortex colored in grey and trabeculae in blue. (J) The indicated bone parameters were quantified. Bone density is bone volume divided by total volume. Trabeculae volume and thickness are normalized to the total inside volume and total bone thickness respectively. (J) A one-way ANOVA was used to test for differences between the doses, followed by post-hoc test for trend. Results are represented as mean ± SEM, dots represent the individual mice. n.s. = not significant, *p<0.05, **p<0.01, ***p<0.001.

Serum GDF15 levels remained stable over time and dependent on the initial EEV dose (**Figure S2B**), indicating a robust *in vivo* overexpression. Furthermore, we observed a dose-dependent decrease in body weight over time, consistent with GDF15’s described role in body weight control (**Figure 2B**)^15,16^. Weight loss caused by both 1µg and 5µg EEV doses overlapped, suggesting receptor saturation. Therefore, we pooled these groups in further read-outs. GDF15-EEV-induced weight loss was confirmed by weighing the visceral adipose tissue depot (VAT) (**Figure 2C**). Together, these data demonstrate that GDF15-EEV as a robust and efficient tool to study the *in vivo* impact of chronically elevated GDF15 levels.

As we observed serum GDF15 levels to be increased in human arthritis, we initially tested the hypothesis that GDF15 itself acts as a proinflammatory mediator. However, GDF15-EEV did not induce clinical signs of inflammation (e.g., joint swelling, skin flaking) or pain (e.g., hunched back, semi-closed eyes) at any dose, with mice appearing otherwise healthy (**Figure 2D**). We extended this analysis using two complementary approaches. First, we evaluated signs of inflammation in the intestine and joint by histopathology. We saw no signs of intestine or joint inflammation in mice treated with GDF15-EEV (**Figure 2E**). In addition, we explored whether GDF15 could alter immune cell composition by in-depth flow cytometry of the spleen, mesenteric lymph nodes and intestine lamina propria. Here we found no differences in immune cell frequencies or distribution with GDF15 overexpression (**Figure S2C-D**). These data indicate that GDF15 itself does not cause inflammation, despite being induced by inflammation.

To further understand the biological role of GDF15 we performed bulk RNAseq on an array of organs. The tissues analyzed included the joint, intestine, liver (the tissue in which the EEV resides) and VAT (known to be affected by GDF15). Globally, there were no GDF15-induced changes in inflammatory genes such as *Nfkb1* and *Stat3* in any of the tissues, nor upregulation of liver-specific inflammation-associated transcription factors (**Figure S2E-F**). *Gdf15* itself was found to be a DEG (Differentially Expressed Gene) in the liver, validating its overexpression (**Figure S2G**). Also in the liver, we found GDF15 to regulate pathways involved in lipid metabolism (**Figure 2F**). In VAT we only found downregulated pathways, involved in angiogenesis and thermogenesis (**Figure 2G**), which reflects fat deposit shrinking. Contrary to the large number of DEG in the liver and VAT, both the intestine and joint showed only a limited amount of DEGs (**Figure S2H**), confirming the lack of pathological changes in these tissues. Collectively, these data confirm that GDF15 induces metabolic changes but found no evidence for it inducing inflammation.

Given the observed association between serum GDF15 levels, SNPs, and bone density in humans, we evaluated bone homeostasis upon GDF15 overexpression. In our RNAseq dataset from the joints, we noticed a dose dependent increase of osteoclast-associated genes, the major bone eroding cell type (**Figure 2H**). We therefore performed µCT (micro-computed tomography) scans of the proximal tibia (**Figure S2I**) and calcaneus (heel bone) (**Figure 2I-J**) to assess extra-articular and peri-articular bone loss respectively. Interestingly, we observed a dose-dependent reduction in total bone density, cortical, and trabecular bone. Thus, GDF15 induces systemic bone loss independently of inflammation.

### GDF15-induced bone loss is GFRAL-dependent

GFRAL is the only known receptor for GDF15 and thus is likely the route through which GDF15 mediates bone loss. To confirm whether GDF15-GFRAL binding is responsible for the observed bone loss, we generated GFRAL-deficient mice (**Figure S3A**), to which we administered GDF15-EEV (**Figure 3A**). As anticipated, we observed a complete protection against GDF15-induced body weight and fat loss in GFRAL-KO mice (**Figure 3B-C**). Next, we assessed the tibia using µCT (**Figure 3D**), which showed that GFRAL-KO mice are protected against GDF15-induced trabecular but not cortical bone loss. We also measured bone strength using a three-point femur bending assay. This indicated that GFRAL deficiency protects against GDF15-induced weakness of the bone (**Figure 3E**).

**Figure 3.**
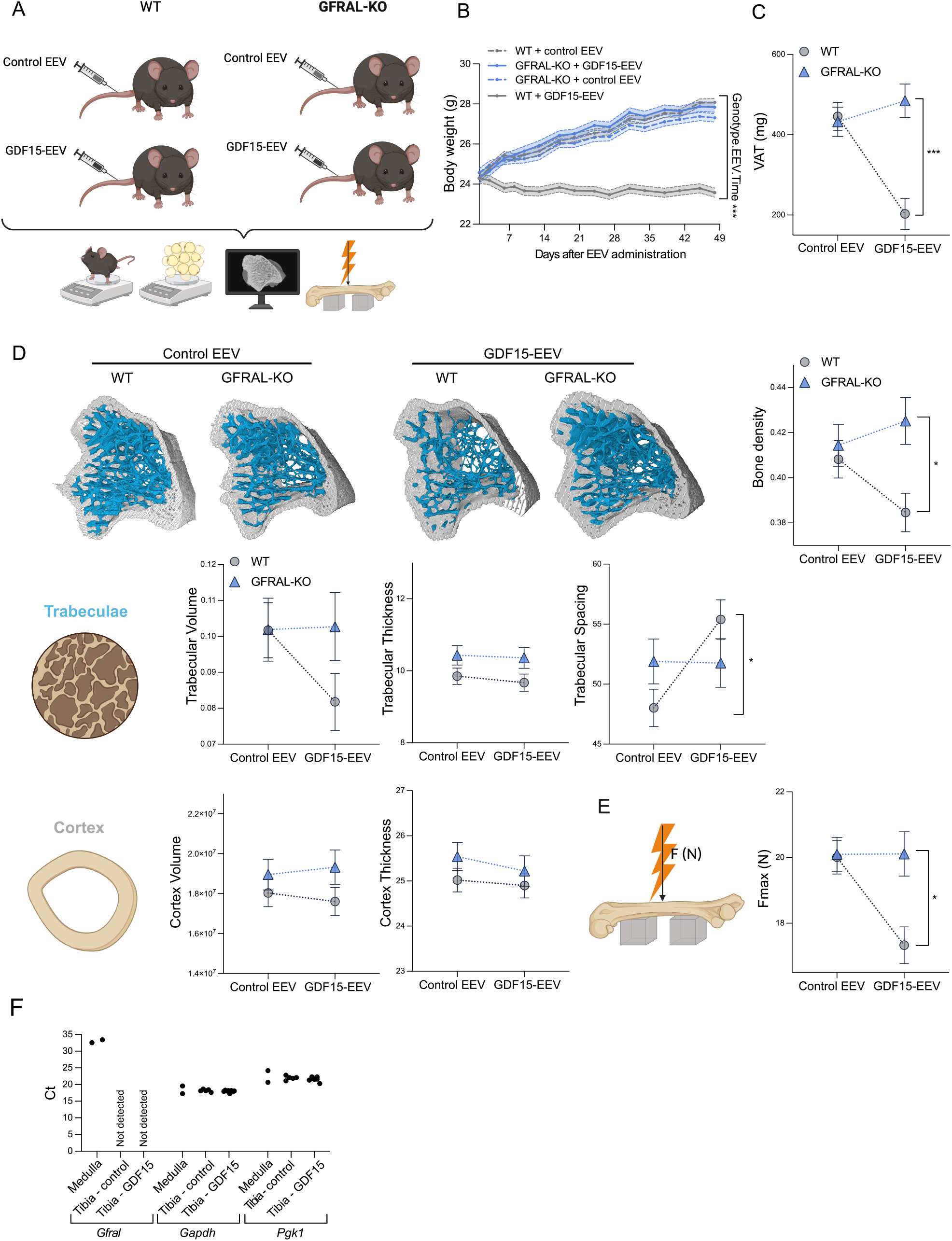
GDF15-induced bone loss is GFRAL-dependent. (A) Schematic overview of the GDF15-EEV GFRAL-KO experiment (n=7-11). GFRAL-KO mice and WT littermates were administered GDF15-EEV or control EEV and monitored for seven weeks. Upon sacrifice, indicated tissues were collected. (B) Mouse body weight over time. Statistics indicate the interaction between the Genotype (KO or WT) and EEV (GDF15 or control) over time as determined by repeated measurements analysis. Body weights are adjusted to baseline body weight. (C) VAT (Visceral Adipose Tissue) weight. (D) Representative µCT images of the tibia. The indicated bone parameters were quantified. (E) Femoral strength assessed by three-point bending test. The Fmax (maximum force) is the load right before the bone fractures. (C-E) A two-way ANOVA was used to test for interactions between Genotype (KO or WT) and EEV (GDF15 or control). (F) qPCR performed on medulla (hindbrain) of WT mice and tibia of mice treated with control or GDF15-EEV. Ct values of the gene of interest *Gfral* and reference genes *Gapdh* and *Pgk1* are shown. Each datapoint represents an individual mouse. Results are represented as mean ± SEM. Significances of the interactions are indicated as: *p<0.05, ***p<0.001.

*Gfral* expression has only been reported in the medulla of the hindbrain. Given we observe that GDF15 affect bones far from the brain, we first ruled out GFRAL expression in bone. While we could detect *Gfral* in the medulla, we could not detect *Gfral* in bone, both at steady-state conditions and upon GDF15 overexpression (**Figure 3F**). Thus, we conclude that GDF15 mediates bone loss through GFRAL, which is not present in the bone, suggesting the existence of a novel brain-bone axis.

### GDF15-induced bone loss occurs independently from weight loss

The GDF15-GFRAL axis is responsible for both body weight and bone loss. Therefore, we investigated the possibility that the observed bone loss is a direct consequence of the reduced caloric intake and/or consequential weight loss. To this end, we performed a pair-feeding experiment. Here, mice were treated with either GDF15-EEV or control EEV. We weighed the amount of food consumed by the EEV-injected groups daily and fed only this amount of food to mice without EEV injection (**Figure 4A**). As previously reported^15,16^, the magnitude of weight loss by elevated GDF15 was replicated completely by caloric restriction (**Figure 4B**). This confirms that reduced food intake is responsible for weight loss caused by GDF15. VAT weight mimicked the whole-body weight measurements (**Figure 4C**). By contrast, bone density was not dependent on caloric restriction: the GDF15-EEV-treated mice had lower total bone density and trabecular bone volume. This was not observed for their EEV-free, pair-fed counterparts (**Figure 4D**). Furthermore, femurs of GDF15-EEV-treated mice require less force to break than their pair-fed counterparts (**Figure 4E**). Thus, GDF15-mediated reduced caloric intake and subsequent weight loss is not responsible for trabecular bone loss.

**Figure 4.**
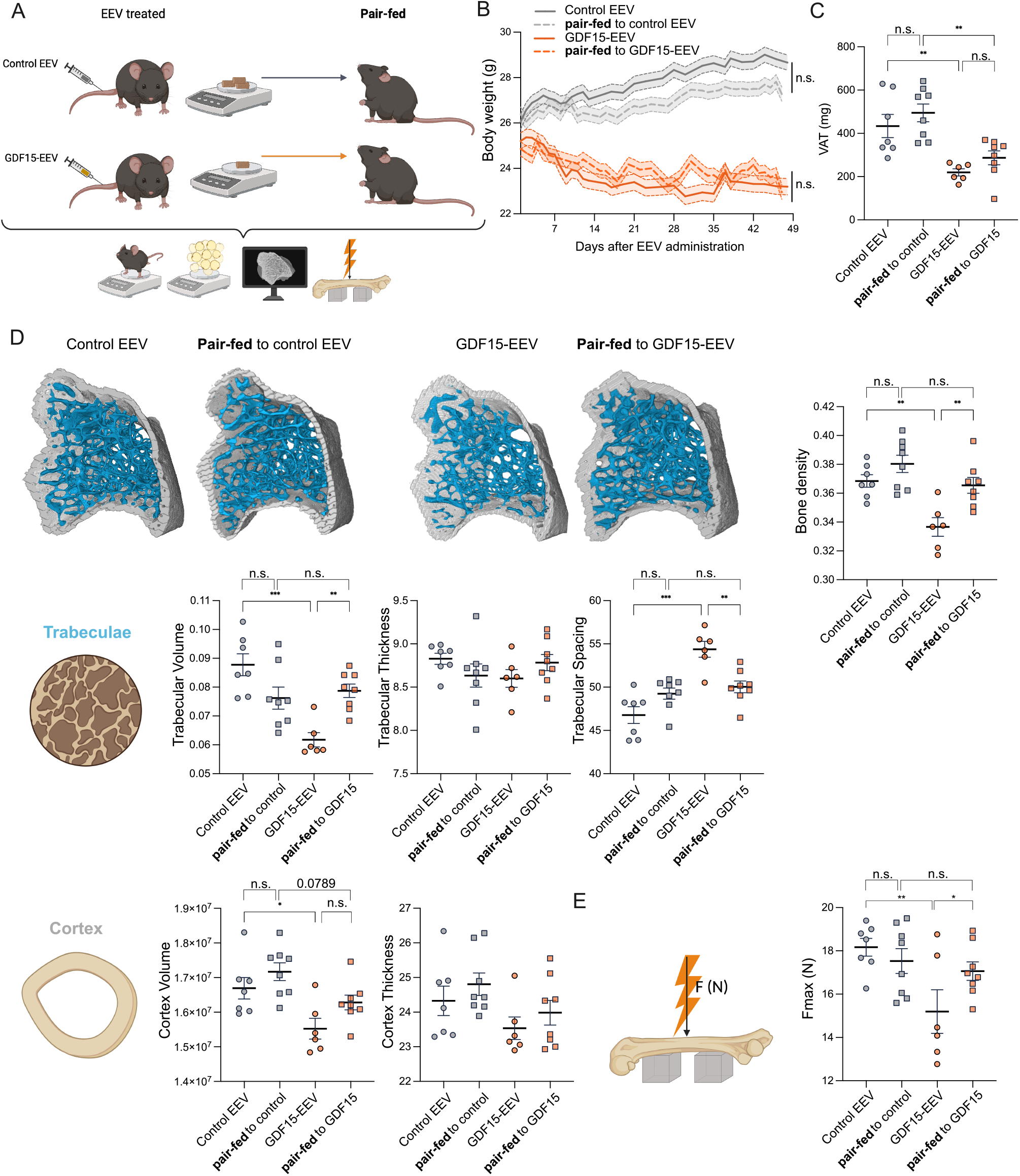
GDF15-induced bone loss occurs independent from weight loss. (A) Schematic overview of the GDF15-EEV pair-feeding experiment (n=6-8). WT mice were treated with control EEV or GDF15-EEV and their food intake was measured daily. An equivalent amount of food was fed to non-EEV injected (pair-fed) WT mice. (B) A repeated measurements analysis was performed to assess overall changes in body weight over time between the groups, represented by the interaction between the EEV-groups, their respective pair-fed groups and time, followed by pairwise contrast testing. Body weights were adjusted for differences in baseline body weight. (C) VAT (Visceral Adipose Tissue) weight. (D) Representative images of µCT of tibia. The indicated bone parameters were quantified. (E) Femoral strength assessed by three-point bending test. (C-E) A one-way ANOVA was used to test for differences between the four groups, followed by post-hoc Šídák’s multiple comparisons test. Results are represented as mean ± SEM. Dots represent the individual mice. Significance of the pairwise comparisons are indicated as: *p<0.05, **p<0.01, *** p<0.001.

### GDF15 is not required for steady-state and non-inflammatory bone homeostasis

As overexpression of GDF15 causes bone loss, we next addressed the question as to whether GDF15 deficiency interferes with steady-state bone homeostasis. Given we observed a gene expression profile of osteoclast activation with GDF15, we first aimed to determine if GDF15 deficiency resulted in a defect in osteoclast development or function. Thus, we cultured osteoclasts from GDF15-KO mice and WT littermates, where we found no difference in the number of multinucleated cells nor in bone resorption (**Figure S3B-C**). Next, we performed µCT bone analysis on adult GDF15-KO and GFRAL-KO mice. We saw no spontaneous bone phenotype of either mouse line (**Figure S3D,F**). Interestingly, these mice also do not show any spontaneous weight phenotype (**Figure S3E,G**). Together, this shows that GDF15 is not required for osteoclast generation, bone homeostasis or body weight in steady-state.

To assess whether GDF15 acts as a universal mediator of bone loss, we initially examined its role in ovariectomy (OVX), a model of post-menopausal osteoporosis^27^. OVX surgery significantly increased body weight compared to the sham intervention, without an effect of GDF15 deficiency (**Figure S4A**). Furthermore, tibia µCT analysis showed that GDF15-KO mice are not protected against bone loss, which was confirmed by femur bending assays (**Figure S4B**). Finally, serum GDF15 levels were unaffected by OVX surgery in WT mice (**Figure S4C**).

In parallel we also examined whether GDF15 mediates age-induced bone loss in view of the age-dependent increase of GDF15^28^. We evaluated 15-month-old male GDF15-KO and WT littermates, which showed no differences in body weight (**Figure S4D**). Likewise, tibia µCT scans showed that overall bone density, cortical and trabecular bone loss were unchanged (**Figure S4E**). Collectively, these findings indicate that GDF15 does not act as an active contributor to bone loss in non-inflammatory conditions.

### The GDF15-GFRAL axis controls IL-23-induced trabecular bone loss, but not inflammation severity

We have thus far shown that GDF15 is increased in several forms of inflammatory arthritis and that GDF15 mediates bone loss but not inflammation. We therefore reasoned that GDF15 could be a critical mediator of inflammation-induced bone loss. To address this, we initially searched for upstream proinflammatory cytokines in the context of arthritis that could trigger GDF15 production. Potential candidates include TNF, IL-17 and IL-23, for which targeted biologicals are currently used for treating RA, SpA, psoriasis, and inflammatory bowel diseases. Therefore, we measured serum GDF15 levels in well-established cytokine-driven preclinical arthritis models and compared this induction to the GDF15-inducing high-fat diet (HFD). Whereas we did not observe significantly increased levels in the TNF^emARE^ arthritis model, a model of SpA^29^, we found that GDF15 levels in the IL-23 EEV model were doubled compared to controls, as it does in HFD (**Figure 5A**). This suggests that IL-23, but not TNF, may induce GDF15 *in vivo*. We therefore focused on the potential functional consequences of the GDF15-GFRAL axis in an IL-23-induced inflammation model.

**Figure 5.**
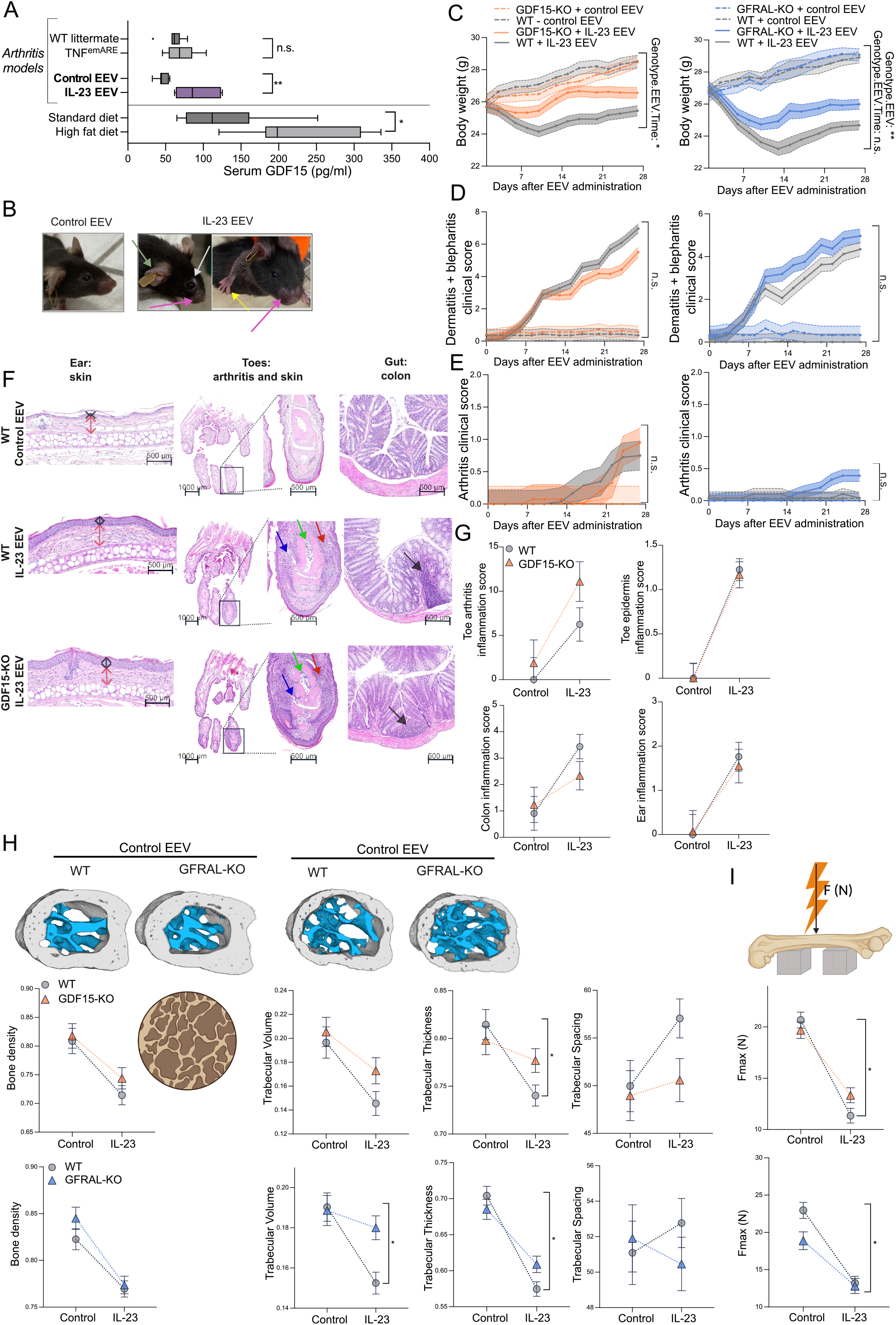
The GDF15-GFRAL axis controls IL-23-induced trabecular bone loss, but not inflammation severity. (A) Serum GDF15 levels of C57Bl/6J mice in cytokine-driven arthritis models and metabolic disease. An unpaired t-test was used to test the differences between disease conditions. (B) Representative images of clinical manifestations of IL-23 EEV administration in WT mice. White arrow shows blepharitis; pink and green arrow shows dermatitis at the snout and ear respectively. Yellow arrow shows DIP (Distal Interphalangeal Joint) swelling. (C) GDF15-KO (orange) or GFRAL-KO mice (blue) and their respective WT littermates were administered either IL-23 EEV or control EEV (control EEV groups: n=5-6, IL-23 EEV groups: n=8-9) and weighed three times per week. A repeated measurements analysis was performed to assess overall changes over time between the groups, represented by the interaction between the Genotype (KO or WT) and EEV (IL-23 or control) and time. Body weights were adjusted for differences in baseline body weight. (D-E) Clinical scores for psoriasis-like dermatitis and blepharitis (D) and arthritis (E) in GDF15-KO or GFRAL-KO mice. A repeated measurements analysis was performed to assess overall changes over time between the groups. (F-G) Representative H&E-stained microscopy images (F) and quantitative analysis (G) of ear, hind paw DIP joint and colon of control EEV or IL-23 EEV-treated WT and GDF15-KO mice. Ear: black arrow indicates epidermal thickness; red arrow indicates dermal thickness. DIP: red arrow shows soft tissue infiltration, blue arrow shows joint space infiltration, green arrow shows bone marrow edema. Colon: black arrow shows immune cell infiltration. (H) Representative images of µCT of the calcaneus. The indicated bone parameters were quantified. (I) Femoral strength assessed by three-point bending test. (G, H, I) A two-way ANOVA was used to assess the interaction between genotype (KO or WT) and EEV (IL-23 or control). Results are represented as mean ± SEM. Significances of the interactions are indicated as: n.s. = not significant, *p<0.05, **p<0.01.

IL-23 overexpression in mice leads to visual psoriasis and arthritis (**Figure 5B),** and is associated with weight and bone loss^30,31^. When validating this model, we observed stable overexpression of IL-23 over four weeks (**Figure S5A**), during which the mice experienced weight loss and psoriasis-like symptoms (**Figure S5B-C**). µCT analysis showed IL-23-induced of bone loss in the tibia and calcaneus (**Figure S5D**). We therefore reasoned that the IL-23 model is valuable for studying inflammation-induced GDF15. To this end, we injected IL-23 EEV in GDF15-KO and GFRAL-KO mice to explore whether inflammation-induced GDF15 contributes to inflammation and/or bone loss.

First, we assessed body weight. GDF15-KO and GFRAL-KO mice displayed partial protection against weight loss (**Figure 5C**), but had no protection against inflammation-induced VAT weight loss (**Figure S5E**). As adipocytes are known to be directly regulated by IL-17^32^, we believe that IL-23 is a stronger regulator of adiposity than GDF15. These data suggest a potential partial involvement of endogenously increased GDF15 in inflammation-induced weight loss.

Contrary to GDF15’s effect on inflammation-induced weight loss, we did not observe any significant difference in clinical dermatitis, blepharitis and arthritis scores between the genotypes (**Figure 5D-E**). We further investigated inflammation severity by histopathology of the skin, gut, and joints (**Figure 5F-G**, **S5F-G**). We found immune cell infiltrates with IL-23 overexpression in all tissues, regardless of the presence or absence of GDF15 or GFRAL. Together, these data indicate that the GDF15-GFRAL axis does not impact tissue inflammation despite being induced by IL-23.

We then assessed inflammation-induced bone loss by µCT. Intriguingly, GFRAL-KO and GDF15-KO mice were protected against trabecular bone loss, but not overall or cortical bone loss (**Figure 5H**, **S5H**). We observed similar trends in the tibia (**Figure S5I-J**). Importantly, GDF15/GFRAL deficiency was found to provide protection of bone strength as femurs of WT mice lost significantly more strength due to IL-23 overexpression than GDF15/GFRAL-KO mice (**Figure 5I**).

In sum, IL-23 induces tissue inflammation independently of GDF15, but weight and bone loss are GDF15-dependent. While IL-23 induces bone loss in both the cortex and the trabeculae, only trabecular bone loss was found to be mediated by GDF15. Therefore, the cell type responsible for GDF15-mediated bone loss must be present in the bone marrow, likely in a trabeculae-proximal niche, rather than in the periosteal bone surface.

### GDF15 activates MALPs to produce RANKL and M-CSF

Next, we aimed to identify the cell in the bone that responds to GDF15 to mediate bone loss. As *Gfral* is not expressed in the bone, we reasoned that GDF15 mediates bone loss indirectly through a brain-bone axis. Since GDF15 has recently been shown to induce β-adrenergic signaling in adipose tissue and liver via GFRAL^33^ and β-adrenergic agonists are known to induce bone loss^34^, we explored publicly available scRNAseq datasets in the bone for β-adrenergic receptors (βARs)^35–39^.

Close examination of adherent bone marrow cells from 3-month-old mice revealed *Adrb2* (β2AR) to be strongly expressed by a novel population of mesenchymal cells, Marrow Adipogenic Lineage Precursors (MALPs) and some hematopoietic cells, but not by osteoclasts nor osteoblasts (**Figure 6A**)^35^. Other subtypes of βARs were not robustly detected in this dataset (**Figure S6A**). Furthermore, MALPs have recently been shown to be able to induce bone loss by stimulating osteoclasts through RANKL and M-CSF expression^40–43^. Thus, we hypothesized that MALPs could be activated by βARs following GDF15 upregulation.

**Figure 6.**
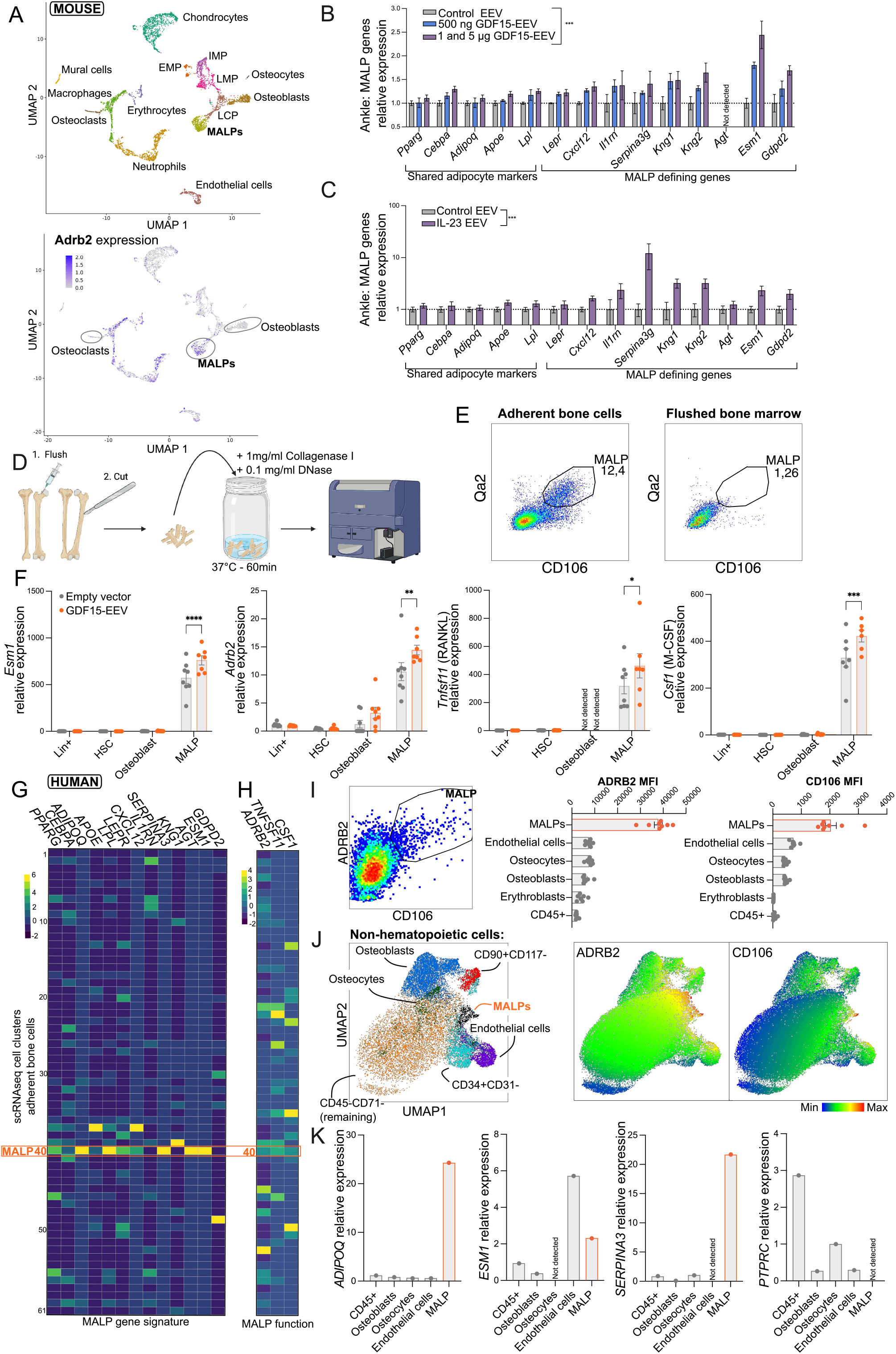
GDF15 activates MALPs to produce RANKL and M-CSF and MALPs are present in human bones. (A) The single-cell dataset (scRNAseq) from 3-month-old mice from Zhong et al.^35^ annotated by clusters and β2-adrenergic receptor (*Adrb2*) expression. EMP = Early Mesenchymal Progenitor; IMP = Intermediate Mesenchymal Progenitor; LMP = Late Mesenchymal Progenitor; LCP = Lineage Committed Progenitor. (B) Bulk RNAseq on whole ankle as discussed in Figure 2 and S2. Counts of each MALP-associated gene is relative to control EEV. A one-way ANOVA was used to test the effect of EEV. (C) Relative Marrow Adipocyte Lineage Precursor (MALP) gene expression determined by qPCR on ankle from mice treated with IL-23 EEV or control EEV, normalized to reference genes *Gapdh* and *Pgk1* and relative to control EEV (n=6). A One-way ANOVA was used. (D) Mouse MALP frequency and activation were assessed using FACS. A schematic of sample processing is shown. First, hind leg bones were cleaned, flushed and cut longitudinally and into small pieces. Bone pieces were digested with collagenase I and DNase for 60min at 37°C, after which cells were stained for FACS. (E) Representative flow cytometry graphs of MALPs in adherent bone cells versus flushed bone marrow. Numbers indicate frequency in the parent population (Lin^-^Sca1^-^CD45^-^CD31^-^PDPN^-^CD90^-^CD34^-^ CD59^-^). (F) Gene expression of Lin+ cells, HSCs, osteoblasts and MALPs of mice overexpressing GDF15 (n=7-8). Relative gene expression of the MALP-defining gene *Esm1,* genes related to osteoclast stimulation (*Tnfsf11* and *Csf1*) and adrenergic receptor *Adrb2* shown. All data were normalized to reference genes *Gapdh* and *Pgk1* and are relative to expression in Lin+ cells. A two-way ANOVA was used, followed by a post-hoc Tukey’s multiple comparison test. (G) scRNAseq on digested human vertebral biopsies. Heatmap of the MALP gene signature by cell cluster. (H) Heatmap showing expression of *RANKL, CSF1* and *ADRB2* by cell cluster. (I) Human MALPs were assessed using FACS on enzymatically digested human knee biopsies. Left, representative flow cytometry image of human MALPs. Right, MFI (Mean Fluorescence Intensity) of selected proteins (n=8). (J) UMAP projection of CD45^-^CD71^-^CD115^-^CD11b^-^ cells with cell populations overlayed (left) and MALP-surface marker expression (right). (K) MALPs and other major cell populations in the bone were sorted and gene expression was determined in two pooled human samples. Expression is normalized to reference gene *GAPDH*. Results are represented as mean ± SEM. Dots represent individual samples. Significances are indicated as: *p<0.05, **p<0.01, ***p<0.001.

To study the putative link between GDF15 and MALPs, we initially examined MALP-associated genes in the bone following GDF15 overexpression *in vivo*. We investigated the “MALP gene signature”, defined by Zhong et al.^43^, a collection of 14 genes that includes shared adipocyte markers, such as *Adipoq* and *Apoe*, but also specific MALP-defining genes. One of these MALP-defining genes is *Esm1,* the sole upregulated DEG in our bulk RNAseq dataset of the joint following 7 weeks of GDF15 overexpression (**Figure S2H**). Close examination of the MALP gene signature in this dataset revealed it to be upregulated in a GDF15 dose-dependent manner (**Figure 6B**). In contrast, osteoblast and hematopoietic cell-associated genes were not upregulated by GDF15, suggesting a unique activation of MALP cells by GDF15 (**Figure S6B**). An additional experiment revealed MALP gene signature upregulation in the tibia to be elevated already after eight days of GDF15 overexpression (**Figure S6C**). Finally, we examined the MALP gene signature in the ankles of mice overexpressing IL-23, and again documented a profound upregulation of MALP genes (**Figure 6C**). Thus, GDF15 appears to regulate the MALP gene signature in the bone in a dose-dependent manner.

To date, MALP cells have only been detected at the cellular level by scRNAseq. We therefore sought to identify MALPs by a high throughput method, specifically flow cytometry. Briefly, we performed *in silico* analysis on the single-cell dataset by Zhong et al.^44^ to identify candidate novel MALP cell surface markers, namely Qa2 and CD106 (*Vcam1*). Using this dataset, we also identified CD59 as a marker to exclude osteoblasts. Tibiae and femurs were flushed to remove non-adherent bone marrow, following which adherent cells were released by enzymatic digestion (**Figure 6D**). Flow cytometry was performed on the adherent bone cells; hematopoietic and non-MALP mesenchymal cells were first gated out following standard bone marrow flow cytometry protocols^45,46^. Within the remaining cells, we found a distinct population of Qa2^+^CD106^+^ cells (**Figure S6D**). Interestingly, these cells were not observed in the non-adherent, flushed bone marrow (**Figure 6E**). To confirm that Qa2^+^CD106^+^ cells are MALPs, we sorted MALPs and other major cell populations. Here we show that only Qa2^+^CD106^+^ cells express MALP genes, thus we call them MALPs (**Figure S6E**).

We next set out to determine how MALPs respond to GDF15. To be able to perform flow cytometry in larger experiments, we adjusted our protocol to include magnetic depletion of the Lin^+^ cells before FACS staining. Upon analysis, we found that four weeks of GDF15 overexpression did not affect the frequency of any target cell types in adherent bone marrow, including MALPs (**Figure S6F**). Next, we performed gene expression analysis on magnetically-sorted Lin+ cells and flow cytometry-sorted hematopoietic stem cells, osteoblasts and MALPs. We found that GDF15 overexpression specifically upregulated the MALP-defining genes *Esm1* and *Kng2* (**Figure 6F**, **Figure S6G**). We also found *Adrb2* to be significantly upregulated in MALPs, suggesting increased β-adrenergic signaling to these cells. Most excitingly, GDF15 overexpression strongly upregulated the expression of *Tnfsf11* (RANKL) and *Csf1* (M-CSF) in MALPs. Thus, GDF15 enhances the ability of MALPs to promote osteoclast differentiation, which in turn causes bone loss.

### MALPs are present in human bones

To date, MALPs have only been identified in mice. We therefore sought to identify MALPs in humans. We first explored scRNAseq data from human vertebral biopsies. Of note, in this dataset there was no additional prior enrichment for mesenchymal cells, resulting in a relatively low resolution for these cell types. Notwithstanding, we found one cluster expressing most of the human orthologues of the murine MALP gene signature (**Figure S7A**, **6G**). Furthermore, we found this cluster to highly express both osteoclast-stimuli *CSF1* and *RANKL* as well as *ADRB2* (**Figure 6H**). These data suggest that in humans MALP-like cells exist, which we aimed to confirm using FACS.

Based on the scRNAseq, we confirmed expression of *VCAM1* in MALPs, but not *Qa2* as this gene has no human ortholog^47^. Thus, we decided to use CD106 (*VCAM1*) and ADRB2 to identify MALPs using flow cytometry on adherent bone cells from human knee biopsies. After gating out cells of known lineages, we detected a distinct population of CD106^+^ADBR2^+^ cells, which we designated as MALPs (**Figure 6I, S7B-C**). Projecting these cells onto UMAPs of non-hematopoietic cells, confirmed MALPs to be distinct from other populations in the bone marrow (**Figure 6J, S7D**). To confirm CD106^+^ADBR2^+^ cells are indeed MALPs, we sorted them and other cell types for qPCR analysis. CD106^+^ADBR2^+^ cells specifically express common adipocyte genes *ADIPOQ* and *CEBPA* as well as MALP-defining genes *ESM1*, *SERPINA3* and *CXCL12*, but not immune-cell associated *PTPRC* (CD45) (**Figure 6K, S7E**). Using these complementary approaches, we conclude that MALPs are conserved across species.

### GDF15 activates MALPs through adrenergic signaling

Lastly, we aimed to identify how MALPs are activated by GDF15. One of the most striking aspects of this interplay, is that the GDF15 receptor GFRAL is uniquely expressed in the brain, suggesting a brain-bone axis. We reasoned that the sympathetic nervous system is most likely responsible, given the increased expression of *Adrb2* on MALPs following GDF15 overexpression. Therefore, we performed two complementary strategies to test this hypothesis (**Figure 7A**). First, we performed chemical sympathectomy using 6-OHDA, which causes degeneration of the terminal ends of the peripheral adrenergic neurons^48^. Both GDF15 and 6-OHDA treatments independently cause weight loss, but do not interact to affect this read-out (**Figure 7B**), as confirmed by comparable VAT weight loss (**Figure 7C**). In striking contrast, 6-OHDA effectively blocked the upregulation of the MALP gene signature by GDF15, indicating that MALP activation is regulated by adrenergic stimulation (**Figure 7D**). Genes associated with other cell types were not affected by 6-OHDA treatment (**Figure S7A**). Thus, GDF15 regulates MALP activation through adrenergic signaling.

**Figure 7.**
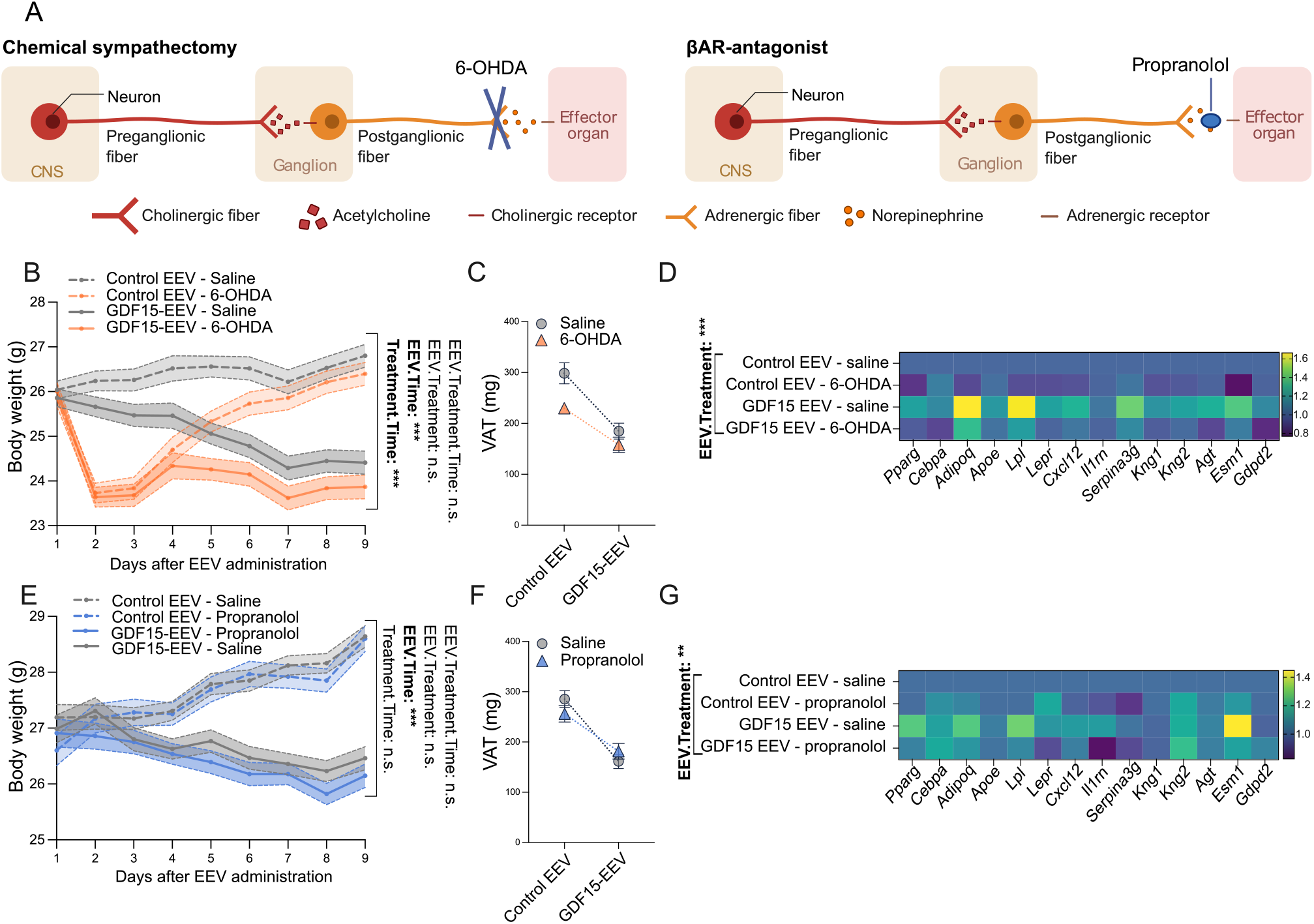
GDF15 activates MALPs through adrenergic signaling. (A) Schematic of the two adrenergic blockade strategies implemented. Left, chemical sympathectomy by 6-OHDA. Right, use of the beta-blocker propranolol. (B, E) Mouse body weight over time of WT mice administered with GDF15-EEV or control EEV and injected with (B) 6-OHDA or (E) propranolol or saline one day after EEV injection (B: n=8-9; E: n=6-8). A repeated measurements analysis was performed to assess overall changes over time in body weight between the groups, represented by the interaction between the EEV (GDF15-EEV or control EEV) and treatment (6-OHDA/propranolol or saline) and time. Body weights were adjusted for differences in baseline body weight. (C, F) VAT (Visceral Adipose Tissue) weight. (D, G) Heatmaps showing the mean gene expression of MALP gene signature in proximal tibia normalized to reference genes *Gadph* and *Pgk1* and relative to the “Control EEV – saline” treated group. (C,D,F,G) A two-way ANOVA was used to assess the significance of the interaction between EEV and treatment. Results are represented as mean ± SEM where applicable. n.s. = not significant, *p<0.05, **p<0.01 ***p<0.001.

In the second approach we used propranolol, a pan-βAR antagonist. Propranolol alone does not affect body and VAT weights, nor does it block the effect of GDF15 (**Figure 7E,F**). In line with our 6-OHDA results, propranolol inhibited GDF15-induced upregulation of the MALP gene signature in the bone (**Figure 7G**). Together, these data indicate that GDF15-induced MALP activation and consequential bone loss is regulated by β-adrenergic signaling.

Altogether, we uncovered a novel role for GDF15 as a mediator of inflammation-associated bone loss. This pathway operates through its receptor GFRAL in the hindbrain, which in turn results in β-adrenergic stimulation of trabeculae-adjacent MALPs. GDF15 increases MALP cell production of RANKL and M-CSF, which controls the differentiation and/or activation of bone-eroding osteoclasts.

## DISCUSSION

Given the intricate relationship between metabolic mediators and inflammation, we anticipated that GDF15 would directly regulate the severity of inflammation in arthritis. Much to our surprise, our experiments revealed the opposite: GDF15 does not appear to modify inflammation in the joints, skin or gut, but controls bone density through a novel brain-bone axis.

The crosstalk between bone and energy metabolism was first described in the early 2000’s. For example, leptin was found to control bone loss through the sympathetic nervous system’s control of osteoblast activity^49,50^. Here, we uncovered a unique link between bone and energy metabolism through GDF15.

Firstly, the link between bone and energy metabolism is usually studied in steady-state conditions and non-inflammatory models of osteoporosis^51^. However, bone appears to be unaffected by GDF15 under these conditions. In stark contrast, during IL-23-driven inflammation, GDF15 mediates trabecular bone loss. This different role of GDF15 in steady state conditions and during inflammation is supported by human data, whereby GDF15 negatively correlates with BMD in RA patients, but not in HCs. Notably, reduced trabecular bone, but not cortical bone, is associated with an increased fracture risk in arthritis patients^52^, underscoring the clinical significance of the trabecular bone. We speculate that the trabecular specificity of GDF15-induced bone loss reflects the close proximity of the target cells to the trabeculae^35^. Thus, it appears that the effect of GDF15 on bone is “specialized” as it primarily affects trabecular bone and does not occur independently of inflammation.

Even more remarkable, GDF15 has no inherent inflammatory properties, yet its removal effectively uncouples inflammation from bone loss. A longstanding prevailing concept states that inflammation-induced bone loss occurs directly through cytokine-mediated activation of osteoclasts^53^. On the other hand, treatment of immune-mediated inflammatory diseases such as RA and PsA by cytokine blockade does not necessarily reverse bone loss despite controlling inflammation^10,54^. Through GDF15, we provide an explanation for this paradox.

Another unexpected aspect of GDF15’s role in brain-bone crosstalk is its ability to regulate weight and bone loss through different mechanisms. This was first demonstrated by our pair-feeding experiment, whereby reduced caloric intake (mimicking GDF15) does not result in the degree of bone loss as GDF15 overexpression. Our chemical sympathectomy and βAR-antagonist experiments confirmed this concept, as we found that the effect of GDF15 on body weight is unaffected by blocking adrenergic signaling, while it blocks the activation of the target cells in the bone.

In this context we identified MALPs as the target cell that links the peripheral nervous system with bone. MALPs, identified in mice as “Lepr-MSC” (Mesenchymal Stem Cell)^38^, “Adipo-CAR” (CXCL12-Abundant Reticular)^39^, “adipo-primed mesenchymal progenitors”^36^ or “pre-adipocytes”^37^, are non-proliferative, non-lipid laden bone marrow adipocyte precursors which have their own unique role in the bone microenvironment. RANKL production by MALPs was shown to be essential for bone homeostasis, which was confirmed by independent research groups^40,41^. Moreover, it was previously assumed that osteoblasts and mature bone marrow adipocytes were the main producers of M-CSF in the bone, however recent evidence shows that MALPs are likely the main source of M-CSF^40,42^. Finally, to the best of our knowledge, we are the first to document the existence of MALPs in human and to show that they express the βAR.

Although MALPs have irrevocably been shown to regulate bone homeostasis, our discovery that they can be regulated by β-adrenergic signaling is novel. β-adrenergic signaling is known to reduce bone mass^55^, and β-blocker use in humans is associated with higher BMD^56^. Older studies stated that osteoblasts are the target cell type for β-adrenergic signaling-induced bone loss^51^. Although we have not formally excluded osteoblasts in GDF15-induced bone loss, recent scRNAseq shows minimal βAR expression on osteoblasts^57,58^. Furthermore, *Tnfsf11*, *Csf1* and *Adrb2* expression was higher in MALPs than osteoblasts, and only MALPs had a further increase in expression of these genes following GDF15 overexpression. MALPs and osteoblasts derive from a common progenitor and both express *Runx2*, which is considered the master transcription factor for osteoblasts^51^. scRNAseq shows that osteoblasts uniquely express *Bglap* (osteocalcin), which we used to confirm these cells in our sorting experiment. Further studies should elucidate the role of MALPs versus osteoblasts in controlling osteoclast activation.

One limitation of this study is that we have not identified the primary source of GDF15. As GDF15 mediates bone loss in inflammatory conditions, we hypothesize that it is expressed at the site of inflammation, potentially by myeloid cells as was described before^59^. Regardless of the source, GDF15 is a circulating protein that exerts its effects in the brain. Future studies should examine the stimuli and cellular source of GDF15 in inflamed tissue. Another limitation of our study is that we have not used targeted approaches to deplete MALPs. MALP-specific KO mice do currently not exist; *Adipoq*-cre conditional depletion used in previous studies targets all adipocytes, including peripheral fat and mature bone marrow adipocytes. It is important to note that in older studies mature bone marrow adipocytes were thought to control bone density^60^. Furthermore, creating MALP-specific KO mice would be very challenging, as MALP-defining genes are also expressed by other cell types.

In summary, we have uncovered a novel brain-bone axis in inflammatory diseases. This axis uncouples inflammation from bone loss. Our study provides evidence that blockade of GDF15 could be an effective strategy to tackle inflammation-associated bone loss. Alternatively, therapeutics from other fields of medicine could be repurposed for the same task: betablockers, known to enhance bone density, could be used to correct arthritis-associated osteoporosis. Finally, as GDF15 is generally upregulated under conditions of inflammation, the knowledge generated herein could be applied to other diseases with upregulated GDF15 in which bone loss is also observed, such as psoriasis and cancer^61,62^.

## ACKNOWLEDGEMENTS

The authors would like to thank all the volunteers to the studies. We thank Amanda Gonçalves of the VIB Imaging core for help with histology images and Tino Hochepied from the VIB Transgenic Mouse core facility for the creation of GFRAL-KO mice. Also, we thank Matthias Jarlborg for sharing IL-23-EEV-treated mouse samples. We thank Robert Inman (University of Toronto) for the IL-23 EEV. Funding was acquired through: D.E., EOS (Excellence of Science, 30480119) and FWO (Fonds Wetenschappelijk Onderzoek – Vlaanderen, G082023N, G067219N, G0C7719N) and European Union IMA (Innovative Medicines Innitiatives) “Health Initiatives in Psoriasis and PsOriatic arthritis ConsoRTium European States” — ‘HIPPOCRATES’). G.S., DFG (Leibniz award) and the European Union (ERC Synergy grant 4D Nanoscope). UGCT Centre of Expertise: The special research fund of the Ghent University (BOF-UGent) is acknowledged for the financial support (BOF.EXP.2017.0007).

## AUTHOR CONTRIBUTIONS

Conceptualization: R.V.d.C., S.L., E.G. and D.E.; Methodology: R.V.d.C., E.G., J.D., I.H. and D.E.; Software: R.V.d.C., D.G., C.V., P.Z., I.H. and D.B.; Formal Analysis: R.V.d.C. and M.V.; Investigation: R.V.d.C., E.G, J.D, I.H., I.J, M.G.R, P.J., S.D.M., L.S., F.S., T.D., J.C., G.P., E.D. and T.M.; Data Curation: R.V.d.C., T.R., L.S., C.V., P.J., Z.L. and E.D; Writing – Original Draft: R.V.d.C.; Writing – Review & Editing: R.V.d.C., E.G., G.S. and D.E.; Visualization: R.V.d.C.; Supervision: E.G., D.E., R.I.D, J.L., L.V., L.V.H., K.R., A.R., G.S. and S.D.; Funding Acquisition: D.E.

## STAR METHODS

### Lead contact

Further information and requests for resources and reagents should be directed to and will be fulfilled by the lead contact, Dirk Elewaut.

## EXPERIMENTAL MODEL AND STUDY PARTICIPANT DETAILS

### Human data

#### GDF15 serum levels detection in HC, RA and SpA patients (cohort 1)

The multiplex assay determining GDF15 serum levels was performed on Belgian subjects, and included 20 healthy controls (HC; 9 male, 11 female), 20 RA patients (4 male, 16 female) and 111 SpA patients (53 male, 58 female) with an expert opinion diagnosis fulfilling the ACR-EULAR for RA and the ASAS classification criteria for SpA at the Rheumatology department, Ghent University Hospital, Belgium. The patients have a respective mean ± SD age of 30.85 ± 6.60, 56.2 ± 14.47 and 35.01 ± 10.54. The SpA patient population consisted of 71 axial SpA patients, 17 peripheral SpA patients and 17 patients with a mixed clinical presentation. Of the SpA patients, 16 had co-morbid psoriasis (either skin or nail). All metadata can be found in **Supplementary table S2**. Ethical approval was obtained from the Medical Ethics Committee of Ghent University (Hospital). All patients gave informed consent.

#### GDF15 serum levels detection in HC and PsA patients (cohort 2)

GDF15 serum levels were determined in German subjects, including 31 HCs (13 male, 18 female) and 110 PsA patients (53 female, 57 male) with a respective mean ± SD age of 39.8 ± 8.42 and 49.37 ± 12.9. Patients were diagnosed using the CASPAR criteria. All metadata can be found in **Supplementary table S3**. Ethical approval was obtained at the Ethical Committee of Erlangen. All patients gave informed consent.

#### GDF15 serum levels detection in PsO patients (cohort 3)

GDF15 serum levels were determined in 29 Belgian PsO patients, diagnosed at the Dermatology department, Ghent University Hospital, Belgium. The patients had a mean ± SD age of 45.03 ± 16.65 and 8 patients were obese. Three patients had a history of joint complaints, but not at the time of blood withdrawal. All metadata can be found in **Supplementary table S4**. Ethical approval was obtained at the Medical Ethics Committee of Ghent University (Hospital). All patients gave informed consent.

#### DEXA and GDF15 serum level correlations in HC and RA patients (cohort 4)

Belgian HC and RA patients with an expert opinion diagnosis fulfilling the ACR-EULAR classification criteria at the Rheumatology department, Ghent University Hospital, Belgium underwent DEXA scanning and serum was acquired. This cohort included 54 HC (20 female, 34 male) with a mean ± SD age of 46.62 ± SD 13.2 and 46 RA patients (33 female, 13 male) with a mean ± SD age of 55.88 ± 12,2. **Supplementary table S5** shows the metadata. Ethical approval was obtained at Medical Ethics Committee of Ghent University (Hospital). All patients gave informed consent.

#### Identification of MALPs in human using single cell RNAseq

Four chronic low backpain patients undergoing spinal fusion surgery at the Balgrist University Hospital, Switzerland, were recruited for bone marrow biopsy acquisition from the unaffected lumbar vertebrae. The patients had a mean ± SD age of 56.8 ± 10.2 and one patient was obese. Three patients were female, one patient was male. Three biopsies were collected from vertebral level L4, one was collected from vertebral level L5. Ethical approval was obtained from the local Ethics Commission and all patients gave informed consent.

#### Identification of MALPs in human using FACS

Biopsies were taken from seven human knee samples received from knee-replacement surgeries of patients with osteoarthritis and one vertebral biopsy. Knee biopsies consist of four femur condyle and three tibia plateau samples. The patients had a mean ± SD age of 61 ± 9.4. Six patients were female, and two were male.

### Mouse data

#### GDF15-EEV titration experiment

9-week-old male C57BL/6J mice were purchased from Janvier Labs (France). Mice were kept in conventional housing conditions with group housing in a 12hr light / 12hr dark cycle. The mice had *ad libitum* access to food and water. The mice were left to acclimatize for 1 week before experiment start at 10 weeks of age. Each experimental group was assigned 4 mice, ensuring that each cage contains mice from each experimental group. Group assignment was random within each cage. The experiment was approved by the Animal Ethics Committee of the Faculty of Medicine and Health Sciences, Ghent University.

#### GDF15-KO and GFRAL-KO steady-state experiment

GDF15 Tm1a mice (C57BL/6J background) were acquired from EUCOMM and subsequently bred with Flpe-deleter and Sox2-cre to create full-KO Tm1d mice on C57BL/6J background. These mice were used before in Maschalidi et al, Nature (2022)^63^. GFRAL-KO mice were created using CRISPR/Cas9 (see methods details). Mice were bred under SPF conditions and group housed in a 12hr light / 12hr dark cycle, having *ad libitum* access to food and water. The mice were moved to conventional conditions upon weening. Male and female KO mice and WT littermates were sacrificed at exactly 14 weeks of age until the predetermined number of mice was reached. The experiment was approved by the Animal Ethics Committee of the Faculty of Medicine and Health Sciences, Ghent University.

#### OVX experiment

Mice were bred under SPF conditions and group housed in a 12hr light / 12hr dark cycle, having *ad libitum* access to food and water. The experiment was performed in conventional conditions. Female GDF15-KO mice and WT littermates were randomly assigned to surgery groups. OVX (ovariectomy) or Sham surgery was performed at 12 weeks of age. To ensure all mice had the same age at the time of surgery, multiple injection groups had to be pooled to reach the predetermined number of mice. The experiment was approved by the Animal Ethics Committee of the Faculty of Medicine and Health Sciences, Ghent University.

#### Ageing experiment

Mice were bred under SPF conditions and group housed in a 12hr light / 12hr dark cycle, having *ad libitum* access to food and water. The mice were moved to conventional conditions upon weening. Male GDF15-KO mice and WT littermates were sacrificed at exactly 15 months of age until the predetermined number of mice was reached. The experiment was approved by the Animal Ethics Committee of the Faculty of Medicine and Health Sciences, Ghent University.

#### GDF15-EEV in GFRAL-KO line

GFRAL-KO mice were bred under SPF conditions. After genotyping, male mice were moved to conventional conditions to perform experiments. The mice were group housed in a 12hr light / 12hr dark cycle and had *ad libitum* access to food and water. Male KO and WT littermates were randomly assigned to the treatment groups. Mice were injected with EEV at 10 weeks of age. To ensure all mice had the same age at the time of injection, injections were staged in groups. The experiment was approved by the Animal Ethics Committee of the Faculty of Medicine and Health Sciences, Ghent University.

#### Pair-feeding experiment

9-week-old male C57BL/6J mice were purchased from Janvier Labs (France). Mice were kept in conventional conditions with individual housing in a 12hr light / 12hr dark cycle. To account for induced stress by individual housing, the mice were given additional enrichment in the form of saw dust and chew bones. The mice were left to acclimatize for one week before the experiment’s start. Mice were allocated into groups based on body weight, ensuring that each group had approximately the same mean baseline body weight. EEV-groups were fed *ad libitum*, pair-fed groups were given limited food each afternoon (see section Methods details). All groups had unlimited access to water. The experiment was approved by the Animal Ethics Committee of the Faculty of Medicine and Health Sciences, Ghent University.

#### TNF^emARE^ mice to determine serum GDF15

TNF^emARE^ mice were recently characterized by Thiran et al. and have arthritis and inflammatory bowel disease^29^. The mice are bred under SPF conditions and subsequently kept in conventional conditions with group housing in a 12hr light / 12hr dark cycle and unlimited access to food and water. Serum GDF15 was determined on samples collected with the purpose of another experiment from male and female mice of 15 weeks of age. The experiment was approved by the Animal Ethics Committee of the Faculty of Medicine and Health Sciences, Ghent University.

#### HFD mice to determine serum GDF15

Serum GDF15 was determined on samples from male, WT C57Bl/6J mice on HFD (High Fat Diet) and SD (Standrad Diet) collected with the purpose of another experiment^64^. The mice were kept in SPF conditions with group housing in a 12hr light / 12hr dark cycle and unlimited access to food and water. Littermate mice were fed either SD (10% kcal fat, Research Diets Inc, Cat. D12450B) or HFD (60% kcal fat, Research Diets Inc, Cat. D12492) from the age of 8 weeks onwards. Serum was collected 12 weeks later (aged 20 weeks). The experiment was approved by the Animal Ethics Committee of the VIB Center for Inflammation Research, Belgium.

#### C57BL/6 IL-23 EEV experiment

9-week-old male C57BL/6J mice were purchased from Janvier Labs (France). Mice were kept in conventional housing conditions with group housing in a 12hr light / 12hr dark cycle and with unlimited access to food and water. The mice were left to acclimatize for one week before the experiment’s start. Accounting for joint pain due to potential arthritis development, additional food was placed on the bedding on the bottom of the cage. Group allocation was random within each cage, ensuring that that each cage has an equal number of mice for each group. The experiment was approved by the Animal Ethics Committee of the Faculty of Medicine and Health Sciences, Ghent University.

#### GDF15-KO and GFRAL-KO IL-23 EEV experiments

Mice were bred under SPF conditions and after genotyping moved to conventional conditions, group housed in a 12h light / 12h dark cycle, having *ad libitum* access to food and water. Accounting for joint pain due to potential arthritis development, additional food was placed on bedding on the bottom of the cage. Male GDF15-KO and WT littermates and GFRAL-KO and WT littermates were randomly assigned to treatment groups and the experiment started at 10-12 weeks of age. To ensure all mice had the same age at the time of injection, multiple injection groups had to be pooled to reach the predetermined number of mice. The experiment was approved by the Animal Ethics Committee of the Faculty of Medicine and Health Sciences, Ghent University.

#### GDF15-EEV short-term experiment to determine MALP gene signature expression by qPCR

9-week-old male C57BL/6J mice were purchased from Janvier Labs (France). Mice were kept in conventional housing conditions with group housing in a 12hr light / 12hr dark cycle and with *ad libitum* access to food and water. The mice were left to acclimatize for one week before experiment start. The mice were assigned to treatment groups based on body weight, ensuring that the mean start body weight of each group was approximately the same and that each cage had the same number of mice of each treatment group. The experiment started at 10 weeks of age. The experiment was approved by the Animal Ethics Committee of the Faculty of Medicine and Health Sciences, Ghent University.

#### Sorting experiment to validate Qa+CD106+ cells as MALPs by qPCR

Three WT C57BL/6J littermate mice which were excess from in-house breeding were assigned to the sorting experiment. Mice were kept in conventional housing conditions with group housing in a 12hr light / 12hr dark cycle and with *ad libitum* access to food and water. The mice were sacrificed at 10 weeks of age. The experiment was approved by the Animal Ethics Committee of the Faculty of Medicine and Health Sciences, Ghent University.

#### GDF15-EEV MALP sort experiment

9-week-old male C57BL/6J mice were purchased from Janvier Labs (France). Mice were kept in conventional housing conditions with group housing in a 12hr light / 12hr dark cycle and with *ad libitum* access to food and water. The mice were left to acclimatize for one week before the experiment’s start. The mice were assigned to treatment groups based on body weight, ensuring that the mean start body weight of each group was approximately the same and that each cage had the same number of mice of each treatment group. The experiment started at 10 weeks of age. The experiment was approved by the Animal Ethics Committee of the Faculty of Medicine and Health Sciences, Ghent University.

#### 6-OHDA experiment

9-week-old male C57BL/6J mice were purchased from Janvier Labs (France). Mice were kept in conventional housing conditions with group housing in a 12hr light / 12hr dark cycle and with *ad libitum* access to food and water. The mice were left to acclimatize for one week before the experiment’s start. The mice were assigned to treatment groups based on body weight, ensuring that the mean start body weight of each group was approximately the same and that each cage had the same number of mice of each treatment group. The experiment started at 10 weeks of age. As 6-OHDA treatment results in severe suffering, we aimed to avoid giving extra stress to the mice by cleaning the cages 2 days before experiment start and not during the beginning of the experiment. Furthermore, extra enrichment was added to the cage by adding extra shelters, saw dust and chew bones. The experiment was approved by the Animal Ethics Committee of the Faculty of Medicine and Health Sciences, Ghent University.

#### Propranolol experiment

9-week-old male C57BL/6J mice were purchased from Janvier Labs (France). Mice were kept in conventional housing conditions with group housing in a 12hr light / 12hr dark cycle and with *ad libitum* access to food and water. The mice were left to acclimatize for one week before the experiment’s start. The mice were assigned to treatment groups based on body weight, ensuring that the mean start body weight of each group was approximately the same and that each cage had about the same number of mice of each treatment group. The experiment started at 10 weeks of age. The experiment was approved by the Animal Ethics Committee of the Faculty of Medicine and Health Sciences, Ghent University.

## METHOD DETAILS

### Serum protein levels

#### Multiplex assay to determine serum GDF15 on human serum (cohort 1)

Serum samples were centrifuged for 10min at 8000g at room temperature (RT) within 2hr of collection. Serum was aliquoted and stored at –80°C. Serum GDF15 levels were determined by an optimized Luminex Bio-Plex immunoassay at the UMC Utrecht Luminex Core Facility (University Medical Center, Utrecht, The Netherlands)^65^.

#### GDF15 ELISA on human serum (cohort 2-4)

Serum samples were centrifuged for 10min at 8000g at RT within 2hr of collection. Serum was aliquoted and stored at –80°C. GDF15 levels were determined using human GDF15 ELISA (R&D, Cat. DY957) with changes to the manufacturer’s protocol: Reagent diluent was replaced by 0.1% casein in PBS and the enzyme was replaced by avidin-HRP (Invitrogen, Cat. 18-4100-51). All samples were diluted 1:5.

#### GDF15 ELISA on mouse serum

Blood was collected either by retro-orbital bleeding or cardiac puncture in micro sample tubes (Sarstedt, Cat. 41.1500.005). Samples were centrifuged at 8000g for 8min at RT and subsequently stored at –20°C. GDF15 levels were determined using a mouse GDF15 ELISA (R&D, Cat. DY6385) with following changes to the manufacturer’s protocol: To block the plate and to dilute the samples 0.1% casein in PBS was used instead of reagent diluent and the enzyme was replaced by avidin-HRP (Invitrogen, Cat. 18-4100-51). Samples were measured undiluted, except for mice injected with GDF15-EEV.

#### IL-23 ELISA on mouse serum

IL-23 levels were determined using a mouse IL-23 ELISA kit (BioLegend, Cat. 433704) following to the manufacturer’s protocol.

### Obtaining GWAS data

The GWAS data was obtained using “gwasrapidd” R package^66^ (online database: https://www.ebi.ac.uk/gwas/home). First, we identified variants for the selected list of genes (*GDF15, GFRAL, ERAP1, IL1R2, IL23R, IL6R, TYK2, HLA-C*) using get_variants() function. Then, we selected variants within 1Mb of the gene body and filtered variants related to genes outside of the original geneset. Function get_associations() was used to get association IDs, p-values, and odds ratio (or_per_copy_number) for every variant, then get_traits() function was used to extract biological traits. The summary graph with several SNPs was plotted using GraphPad Prism software (version 10.1.0). The R software version was 4.2.2 and “gwasrapidd” package version was 0.99.14. For the phenotypes, SNP rs10807491 is specifically associated with heel BMD.

### EEV design, production, and *in vivo* protocols

#### In silico design of GDF15-EEV and IL-23

Mouse *Gdf15* transcript sequence was obtained from NCBI (NM_011819). The protein coding exons were flanked *in silico* by the multiple cloning site (MCS) + 10 adjacent base pairs from the CAGs-MCS EEV (Systems Bio, Cat. EEV600A-1). The *Gdf15* transcript was examined for restriction sites *in silico* (SnapGene) to select for appropriate restriction enzymes to use for cloning. IL-23 EEV was a gift from R. Inman (University of Toronto). It was originally purchased from Systems Bio (Cat. EEV651A-1).

#### EEV cloning

The *Gdf15* sequence was generated at VIB synthetic DNA core facility using a BioXP. To linearize the EEV, NotI (NEB, Cat. R189S) and BsrGI (NEB, Cat. R3575S) restriction enzymes were used. *Gdf15* was inserted into the linearized vector using Gibson assembly (NEB, Cat. E5520S). Next, the EEV was transformed into Stbl2 competent cells by heat shock following the manufacturer’s instructions (ThermoFisher, Cat. 10268019). After serial dilution in SOC media with ampicillin, colonies were grown on agar plates (37°C). Six colonies were picked the next morning, and the culture was expanded in 100ml Luria broth with overnight shaking (200rpm, 37°C). 500µl of transformed Stbl2 cells from overnight culture were cryopreserved in 35% glycerol for future use. A 1.5ml aliquot of Stbl2 cells from overnight culture was used for miniprep plasmid extraction (QIAGEN Plasmid mini kit, Cat. 12123), to check the EEV sequence. The remaining cells were pelleted by centrifugation (4000rpm, 30min, 4C) and stored at –20°C for bulk EEV extraction by endotoxin-free maxiprep (Qiagen EndoFree Maxi Kit, Cat. 12362).

IL-23 EEV was revived by firstly adding a scraping of the cryopreserved Stbl2 expressing IL-23 EEV to 5ml Luria broth and incubating at 37°C for 8hr with shaking. 100ul of the starter culture was then expanded overnight in 100ml Luria broth as above. EEV-containing bacteria were collected by centrifugation as above.

#### GDF15-EEV insert sequencing

After miniprep isolation of EEV, PCR was performed using primers flanking the Gdf15 insert (Fw: GGGCAACGTGCTGGTTATTG, Rev: AGCAGCGTATCCACATAGCG). The PCR product size was first assessed by gel electrophoresis to ensure an insert of the expected size was present (1116bp). Colonies containing *Gdf15* inserts of the correct size were sent for Sanger sequencing (Eurofins) to ensure the correct sequence was inserted. Based on the results, we selected a single GDF15-EEV to use for *in vivo* experiments.

#### EEV injection by hydrodynamic delivery (HDD)

EEV was administered by HDD, in which EEV in PBS at a volume of 10% of the mouse’s body weight is injected by tail vein in less than 10 seconds. For GDF15-EEV, in the titration experiment mice were injected with 100ng, 500ng, 1µg or 5µg. In all following experiments, mice were injected with 1µg. For IL-23, mice were injected with 30ng as in our hands higher concentrations can lead to IL-23 induced morbidity (unpublished). The same amount of control EEV was injected for the respective experiments. The mice were weighed before injection to determine the volume of PBS to be injected. Serum was collected one week after injection to determine GDF15/IL-23 serum levels. Mice were excluded based on failed injections (e.g., insufficient volume of too slow) and/or low serum GDF15 or IL-23 levels as determined by ELISA.

#### GDF15-EEV in vivo clinical scoring

In the GDF15-EEV titration experiment, mouse body weight was measured daily for the first week and then three times per week thereafter. At these timepoints the mice were monitored for general signs of inflammation and pain by two blinded investigators up to seven weeks after EEV injection. The following were assessed for clinical signs of inflammation: joint swelling, general coat condition, skin flakes, ear/paw thickening, mobility and alertness, position of eyes and ears, posture, and stool consistency. As no signs of inflammation were seen in the titration experiment, the following experiments, body weight was determined at following time points: before the injection, one day after the injection, followed by minimum three times per week.

#### IL-23 EEV in vivo clinical scoring

Mouse body weight and clinical score was assessed by two blinded investigators at baseline, one day after the injection, and then three times per week, according to **supplemental table S6**. The mice were sacrificed four weeks after EEV injection.

### Study-specific *in vivo* mouse protocols

#### OVX surgery

GDF15-KO and WT mice were randomly assigned to surgery groups (OVX or Sham surgery). The mice were anesthetized using ketamine (100mg/kg body weight) and xylazine (16mg/kg body weight). After shaving, a dorsolateral incision of 5-10mm was made. The ovary and fallopian tube were brought outside of the abdomen, and the vasculature between ovaries and uterus was tied off. The ovary was removed, and the remaining tissue was placed back. The skin was closed using resorbing thread. The procedure was then repeated on the other ovary. Immediately after surgery, the mice received buprenorphine as analgesia (0.1mg/kg body weight). Ovary removal was confirmed using histology on the removed tissue. Mouse body weight was assessed once weekly. For the sham-surgery mice the procedure was the same except for tying the blood vessels and removal of the ovaries. The mice were euthanized six weeks after surgery.

#### Pair-feeding experiment

WT mice were divided into four groups: 1) Control EEV, 2) Pair-fed to control EEV, 3) GDF15-EEV, 4) Pair-fed to GDF15-EEV. Group 1 and 3 received EEV injections on day 0, and starting from day 1, the remaining food in the food hopper was measured daily. To ensure uniformity, the mean amount of food consumed by the EEV group was given to the corresponding pair-fed mice. Both food intake and body weight were monitored daily, with pair-fed mice receiving their food in the afternoon. At day 6, an unexpected recovery in body weight was observed in the pair-fed to GDF15-EEV group. Concurrently, food intake measurements in the GDF15-EEV group exhibited a sudden increase. Upon closer investigation, it became apparent that the GDF15-EEV mice were chewing on and pulverizing food without actual consumption, thereby affecting the accuracy of food intake measurements. This behavior could be a consequence of stress induced by single housing conditions combined with GDF15 overexpression. Consequently, moving forward, any EEV-treated mice displaying signs of food destruction were excluded from the measurement of mean food consumption. Furthermore, efforts were made to enhance the enrichment within all cages to alleviate stress levels. As a result of these observations and adjustments, body weight measurements taken between days 6 and 10 post-EEV injection for the pair-fed to GDF15-EEV mice were omitted from the statistical analysis. Ultimately, all mice were euthanized seven weeks after the initial EEV injection for further investigation.

#### 6-OHDA experiment

WT mice were divided into four groups: 1) GDF15-EEV + 6-OHDA, 2) GDF15-EEV + saline, 2) Control EEV + 6-OHDA, 4) Control EEV + saline. The mice were injected with EEV on day 0. The mice received a single injection with 6-OHDA (Sigma, Cat. H4381) or saline one day after EEV injection. The 6-OHDA solution was prepared by diluting it in saline with 0.1% ascorbic acid (Sigma, Cat. A34403), kept on ice, and was injected within 30min of dissolving. These specific conditions were applied to avoid breakdown of the product, observable by a color change of the solution which did not occur. 6-OHDA was injected intraperitoneally at a concentration of 200mg/kg. For the “saline” control groups, mice received injections of 200µl of saline with 0.1% ascorbic acid. All mice were weighed daily and ultimately sacrificed on day 9. Longer experiments were not performed as 6-OHDA injury is reversible after approximately two weeks^67^.

#### Propranolol experiment

WT mice were divided into four groups: 1) GDF15-EEV + propranolol, 2) GDF15-EEV + saline, 3) Control EEV + propranolol, 4) Control EEV + saline. The mice were injected with EEV on day 0. From day 1 onwards, the mice received daily intraperitoneal injections with propranolol (Sigma, Cat. P0884) or saline. Propranolol was administered at a dosage of 10mg/kg of body weight and was prepared by diluting it in saline. Daily weight measurements were recorded throughout the study period. It’s noteworthy to mention that some unexpected deaths occurred among the GDF15-EEV + propranolol mice for reasons that remain unknown. Consequently, these mice were excluded from the subsequent analysis. For this reason, we have not performed long-term propranolol treatment. Finally, the remaining mice from all four groups were euthanized on day 9 for further examination.

### Generation of the *Gfral*-KO line (official name: Gfral^em1Irc^)

Gfral^em1Irc^ mice (MGI ID: 7526694) were generated using the CRISPR/Cas9 system. Synthetic Alt-R® CRISPR-Cas9 crRNA (Integrated DNA Technologies) with protospacer sequences 5’ AAAGTTTGTTTACTGTACAG 3’ and 5’ TAACCTGAGTATCCAGGCTT 3’ were duplexed with synthetic Alt-R® CRISPR-Cas9 tracrRNA (Integrated DNA Technologies). cr/tracrRNA duplexes (100ng/µl) were complexed with Cas9-eGFP Nuclease (500ng/µl) (VIB Protein Core). The resulting RNP complexes were electroporated into C57BL/6J zygotes using a Nepa21 electroporator with electrode CUY501P1-1.5 using following electroporation parameters: poring pulse = 40V; length 3.5ms; interval 50ms; No. 4; D. rate 10%; polarity + and transfer pulse = 5V; length 50ms; interval 50ms; No. 5; D. rate 40%; polarity +/-. Electroporated embryos were incubated overnight in Embryomax KSOM medium (Merck, Millipore) in a CO2 incubator at 37°C. The following day, 2-cell embryos were transferred to pseudopregnant B6CBAF1 foster mothers. The resulting pups were screened by PCR over the target region using primers 5’ CCTGGCACTTTGAGTATT 3’ and 5’ TAGCCAGCATTAGACCATT 3’. PCR bands were Sanger sequenced to identify the exact nature of the deletion. Mouse line Gfral^em1Irc^ contains an allele with a deletion of 58bp (chr 9:76112664-76112721) in exon 3 (ENSMUSE00000496764) of the *Gfral* gene (ENSMUSG00000059383). This deletion creates a frameshift resulting in premature stop codons and NMD. All base annotations are according to C57BL/6J genome assembly GRCm39.

### Histopathology

#### Tissue processing

Upon dissection, tissues were fixed in 4% formaldehyde overnight for soft tissues (gut, ear, liver) or 48 hours for hard tissues (ankle, foot). Then, tissues were transferred to 70% ethanol until processing. Hard tissues were decalcified using 5% formic acid for five days. After dehydration and paraffinization, samples were embedded in a paraffin block and cut using a microtome at a thickness of 5µm. Samples were stained using hematoxylin and eosin to assess inflammation.

#### Tissue histopathology scoring

Two investigators assessed the inflammation severity in a blinded manner using the previously reported scoring systems for different anatomical regions^68^. Briefly:

Ear: The ear was used as a skin read-out. The epidermis was scored on a scale of 0 to 3, with the score being based on the number of layers of keratinocytes. Additionally, the dermis was scored from 0 to 4, considering thickness and the extent of immune cell infiltration.

Colon: Assessment of the colon involved scoring based on the severity (ranging from 0 to 3) and extent (also ranging from 0 to 3) of infiltration, as well as evaluating epithelial changes, including crypt elongation (0-3) and goblet cell loss (0-3).

Ankle: In the ankle region, several anatomical regions were taken into account. Samples were scored for Achilles tendonitis (0-2), calcaneus bone marrow edema (0-2), and infiltrate in the synovial-Achilles enthesial complex (0-2). Further evaluations included scoring the talus-tibia-calcaneus region (0-2) and assessing the areas around the cuboidal joints, such as the synovium (0-2), fat pad (0-2), and joint space (0-2).

Toes (hind foot): For each joint in each toe, periarticular inflammation (0-3) and joint space infiltration (0-2) was assessed. For each toe, overall bone marrow edema was scored (0-3). For each sample, epidermis thickness was evaluated based on the number of cell layers at the base of the toes (0-3).

### Bone assays

#### µCT

Scanning: Mouse hind paws where carefully dissected and fixed using 4% formaldehyde for 48hr, after which the samples were transferred to 70% ethanol awaiting scanning. The scans were performed at the Centre for X-ray Tomography of the Ghent University (UGCT, www.ugct.ugent.be) using the HECTOR micro-CT scanner^69^. This system makes use of an X-RAY WorX directional type X-ray tube and was updated with a Varex XRD 4343 flat-panel detector with CsI scintillator, measuring 43×43cm^2^ and having 2880×2880 pixels with a pixel pitch of 150μm. The scans were performed using a tube voltage of 130kV, a tube power of 10W and no beam filtering was applied. During each scan 2001 projections with an exposure time of 1 second per projection were taken covering a full sample rotation of 360^0^. During scanning, the tibiae were kept in a closed tube containing also a few drops of 70% ethanol to prevent tissue dehydration. Before the reconstruction of each scan the projection images were clipped to 2200×2200 pixels. The 3D reconstruction was performed using Octopus, a reconstruction software package developed at UGCT^70^, yielding a 10.6 gigavoxel reconstructed volume with an isotropic voxel size of 4μm.

Analysis: µCT data analysis was performed on tibia and calcaneus as described in detail in Gilis *et al.*, Arthritis Rheumatology (2019)^71^ and in Cambré *et al*., Nature Communications (2018)^72^ respectively. Briefly, to quantify bone density parameters such as bone volume over total volume and cortex and trabeculae thickness, an in-house developed script was used in the ImageJ software. Here, the bone structures were automatically classified into cortex and trabeculae using an algorithm similar to that of Buie *et al*.^73^. Bone regions were quantified by measuring both the average thickness of these structures (using the Thickness plugin from BoneJ) as well as their entire volume. The “bone density” measure is the ratio of the “bone volume” over the “total volume”. The “trabecular spacing” measure is the average thickness of the non-bone volume between the trabeculae. “Trabecular volume” was normalized by dividing by the “inside volume”, which is calculated by subtracting the “cortical volume” (including cortical pores) from the “total volume”. In calcaneus, “trabecular thickness” was normalized to “total bone thickness”.

#### Femur bending assay

The femurs were carefully excised at the hip and knee joints. All surrounding tissue was removed, and the femurs were wrapped in a PBS-soaked gauze to prevent dehydration. Samples were stored at – 20°C up until analysis. The three-point femur bending test was performed using the LRXplus (Lloyd Instruments, USA) universal testing machine located at the Department of Human Structure and Repair, Ghent University. Here, a loading point is strategically positioned on the mid-diaphysis of the femur and is gradually moved downward, applying increasing force and displacement. The resulting maximum force, measured in Newtons (load cell 100N), signifies the load applied immediately before the femur fractures.

#### In vitro osteoclastogenesis

Bone marrow cells were flushed from mouse femurs and tibiae and subsequently cultured in α-minimum essential medium (α-MEM) supplemented with 10% fetal calf serum, 10 units/ml penicillin, 10mg/ml streptomycin, 2mM glutaMAX and 30ng/ml M-CSF (obtained from VIB protein core facility). After overnight incubation, the non-adherent fraction was seeded at 10^6^ cells/well in 24-well plates in medium containing 30ng/ml M-CSF and 10ng/ml RANKL (R&D Systems, Cat. 452-TEC-019). The medium was replaced with supplemented M-CSF and RANKL after 48hr. Four days after start of the culture, osteoclasts were formed and TRAP staining was carried out according to the manufacturer’s instructions (Sigma-Aldrich, Cat. 386A-1KT). Cell counts were determined by capturing three images in the same region of each well, and these images were then analyzed using ImageJ to count the number of cells. For functional analysis, the above-described culture was conducted on coated bone resorption plates (Bio-Connect, Cat. CSR-BRA-24), which emit a fluorescent signal when resorption occurs. Bone resorption was measured according to the manufacturer’s instructions. As a negative control, RANKL was omitted from the culture medium.

### RNA isolation

For tissues: Upon sacrifice of the mice, tissues were collected and placed in RNA Protect (Qiagen, Cat. 76106). For tibia, the proximal 1/3^rd^ of the cleaned bone is used. RNA was extracted by mixing of the tissue in TRIZol (Life Technologies, Cat. 15596-026) followed by the Qiagen RNeasy micro kit (Cat. 74004) according to the manufacturer’s instructions. For adipose tissue (VAT), an extra centrifuge step (10min, 4°C, full speed) was added after tissue mixing, to eliminate the resulting top fatty layer.

For sorted cells: Cells were centrifuged, and the pellet was lysed using RLT buffer (mouse cells) or QIAZol (human cells) (Qiagen, Cat. 79306). RNA was extracted using the Qiagen RNeasy micro kit.

### Mouse bulk RNAseq

#### Sequencing

RNA concentration and purity were determined spectrophotometrically using the Nanodrop ND-8000 (Nanodrop Technologies) and RNA integrity was assessed using a Fragment Analyzer (Agilent). Per sample, an amount of 500ng of total RNA was used as input. Using the Illumina TruSeq® Stranded mRNA Sample Prep Kit (protocol version: Part # 1000000040498 v00 – October 2017) poly-A containing mRNA molecules were purified from the total RNA input using poly-T oligo-attached magnetic beads. In a reverse transcription reaction using random primers, RNA was converted into first strand cDNA and subsequently converted into double-stranded cDNA in a second strand cDNA synthesis reaction using DNA PolymeraseI and RNAse H. The cDNA fragments were extended with a single ‘A’ base to the 3’ ends of the blunt-ended cDNA fragments after which multiple indexing adapters were ligated introducing different barcodes for each sample. Finally, enrichment PCR was carried out to enrich those DNA fragments that have adapter molecules on both ends and to amplify the amount of DNA in the library. Sequence-libraries of each sample were equimolarly pooled and sequenced on Illumina NovaSeq 6000 (v1 kit, 100 cycles, Single Reads) at the VIB Nucleomics Core (www.nucleomics.be).

#### Data processing

The quality of raw sequencing reads from 48 bulk RNA-seq samples, categorized into four subsets visceral adipose tissue (VAT, n=12), ankle (n=12), small intestine (SI, n=12), and liver (n=12), was evaluated using fastQC v0.11.9. Subsequently, untrimmed reads were aligned to the mouse reference genome GRCm38 utilizing HISAT2 v2.2.0^74^. The read counts were generated using featureCounts v2.0.0^75^. For each individual subset, differential expression analysis was performed using DESeq2 v1.32.0^76^ in R v4.1.0. Filtering was lenient, excluding genes with fewer than 10 counts. The DESeq() function, utilizing default parameters, was employed for read count normalization, dispersion estimation, and linear model fitting. Principal component analysis was utilized to identify potential batch effects and outliers. Notably, outlier identification led us to identify one outlier in subset VAT and one in the ankle subset. To mitigate the impact of lowly expressed genes and high variability, we applied the apeglm shrinkage estimator to the log fold changes^77^. Significance levels were adjusted for multiple testing using the Benjamini-Hochberg (BH) method. Genes were considered differentially expressed if they passed the commonly used thresholds of an adjusted p-value <0.05 and an absolute log2 fold change >1.

#### Pathway analysis

To identify enriched Gene-Ontology biological processes (GO-BP), overrepresentation analysis (ORA) was carried out by enrichGO, a function of the R package clusterProfiler v4.0.5^78^. Prior to the analysis, gene symbols were converted to Entrez ID’s in R using AnnotationDbi v1.51.1 and the annotation package org.Mm.eg.db v3.13.0. A custom background gene list was provided containing all genes of the filtered count matrix. The analysis was performed on up– and downregulated genes separately, adjusting the p-values of the enrichment result for multiple testing with the BH method. GO-terms with an adjusted p-value < 0.05 were considered significantly enriched.

### Human single cell RNAseq

#### Sample acquisition and generation of the single cell suspension

From chronic low back pain patients undergoing spinal fusion surgery at the Balgrist University Hospital, Switzerland, a bone marrow biopsy was collected with a Jamshidi bone marrow biopsy needle (HS HOSPITAL SERVICE S.P.A., 8G x 100mm) using the pedicle screw trajectories prior to screw insertion. The vertebral body from which the biopsy was collected did not show any sign of abnormality on magnetic resonance images. MR images were graded by a radiologist with >14 years overall experience and with >7 years in musculoskeletal radiology (NAFA) based on available sagittal T1-weighted (T1w), T2w, Short Tau Inversion Recovery (STIR), and coronal T2w sequences. The mean difference from MRI acquisition to date of surgery was 35 ± 45.99 days. In the operating room, biopsies were immediately transferred to Hanks’ Balanced Salt solution (HBSS) (Sigma-Aldrich). To isolate cells, biopsies were digested for 40min at 37°C in digestion solution (0.05% Collagenase P (Sigma-Aldrich), 100μg/ml Liberase (Sigma-Aldrich), 100μg/ml DNAse I (Sigma-Aldrich) in HBSS, gently flushed with digestion solution, digested for another 20min at 37°C, and filtered through a 70μM filter. Red blood cells were lysed with ACK lysis buffer and viability was determined with Trypan blue (ThermoFisher Scientific). Cell viability had to be > 70% in order to proceed. Cells positive for CD45 and CD66b (Biolegend) and Zombie Aqua (Biolegend) (dead cells) were removed by FACS (FACSAria™ Fusion, BD).

#### Single cell sequencing

A LUNA-FX7 Automated Cell Counter (Logos) was used to determine the cell viability and concentration. Approximately 16,500 cells per sample were loaded onto the 10× Chip G for the experiment, and it was expected that around 10,000 cells could be recovered per library preparation. Cells were then combined with a master mix that contains reverse transcription reagents. The single cell 3’ v3.1 gel beads, carrying the Illumina TruSeq Read1, a 16bp 10× barcode, a 12bp UMI and a poly-dT primer, were loaded onto the chip along with oil for the emulsion reaction. The Chromium X system partitions the cells into nanolitre-scale gel beads in emulsion (GEMs), where the process of reverse-transcription (RT) takes place. All cDNAs within a GEM share a common barcode. After the RT reaction, the GEMs were broken and the full length cDNAs was captured by MyOne SILANE Dynabeads and then amplified with 11 cycles. The amplified cDNA was cleaned up with SPRI beads. After purification, the cDNAs were qualitatively and quantitatively analysed using an Agilent 4200 TapeStation High Sensitivity D5000 ScreenTape. The cDNA was then enzymatically sheared and in the meantime the end of fragments was repaired and A-tailed. Subsequently, sequencing libraries were constructed using the double-sided size selected cDNA fragments. This involved adapter ligation, a sample index PCR with 13 cycles, and SPRI bead clean-ups. The sample index PCR incorporated a unique dual index for sample multiplexing during sequencing. The final libraries contained P5 and P7 primers required for Illumina bridge amplification. Sequencing was performed on an Illumina sequencing platform (Novaseq 6000) using paired-end 28+90bp sequencing. One end of the sequencing read generated cell-specific, barcoded sequences and unique molecular identifier (UMI), while the other end captured the sequence of the expressed poly-A tailed mRNA. The sequencing was carried out using two full FP flow cells to achieve an approximate read count of 50,000 per cell.

#### Data processing

Preprocessing, quality control, and normalization: First, barcodes were processed, genes were aligned to the human genome (Homo_sapiens/GENCODE/GRCh38.p13/Annotation/Release_42-2023-01-30), and a count matrix was generated using the CellRanger toolkit (10× Genomics, version 4.0). Subsequent analyses were performed in R studio (R version 4.2.2). Quality of cells was controlled and cells with mitochondrial content <50% and >500 counts per cell were retained (*scater,* version 1.26.1). Genes with 0 counts across all cells were removed. Data was normalized with sample as batch factor (*batchelor,* version 1.14.1).

Dimensionality reduction, clustering, integration: The top 57% (10695 out of 18651) highly variable genes (HVG) were identified (*scran,* version 1.26.2). Data was integrated on these HVGs, with number of nearest neighbors 30, and without cosine normalization with fastMNN function (batchelor*, version 1.14.1*). Clustering was performed on the corrected low-dimensional coordinates by building a shared-nearest-neighbor graph with 30 neighbors and “rank” weighting scheme (*scran*, *version 1.26.2*) and cluster detection with the Leiden algorithm using resolution of 0.75 (*igraph,* version 1.5.1).

Other parameters were left as default. Grouped heatmaps were created with plotGroupHeatmap, with center and scale, but without clustering rows or columns (scater, version 1.26.1). Viridis color scheme was used (viridisLite, version 0.4.2).

### qPCR protocol and analysis

#### Mouse

After nanodrop measurement to determine concentration and quality of RNA, cDNA was synthesized with the QuantiTect Reverse Transcription kit (Qiagen, Cat. 205314). Gene expression was determined using SYBR Green (GC biotech, Cat. QT650-05) and a LightCycler 480 (Roche) according to the manufacturer’s instructions. Analysis was performed using qbase+ (CellCarta).

Primers were designed in-house. We utilized NCBI’s primer design tool to generate primers spanning exon junctions if possible and where applicable covering all splice variants. The primers for *Bglap* cross-react with *Bglap2* and *Bglap3*, as *Bglap-*specific primers were not possible to design. All paralogues are specifically expressed by osteoblasts. Primers were ordered from IDT and their efficiency and specificity was determined by performing a dilution series with melt curve. To identify optimal reference genes, we evaluated eight candidates, and their suitability was assessed using the geNorm tool available in qbase. Ultimately, *Gapdh* and *Pgk1* emerged as the most reliable reference genes for our study. All the primer sequences used can be found in **Supplemental Table S7**.

#### Human

After nanodrop measurement to determine the concentration and quality of RNA, cDNA was synthesized with the QuantiTect Reverse Transcription kit. Primers for *ESM1, PTPRC, CEBPA,* and *ADIPOQ* were designed in-house. The expression of these genes was determined using SYBR Green (GC biotech, Cat. QT650-05). Primers for *CXCL12*, *SERPINA3*, and *GAPDH* were ordered from Termofisher. These were analyzed using the Fast Advanced Master Mix (Taqman, Thermofisher). Primer information can be found in **Supplemental Table S7**.

### Reconstruction of the MALP dataset acquired by Zhong et al^35^

We utilized the raw UMI count text file of 3-month-old mice from the Gene Expression Omnibus (GEO) series GSE145477 to reconstruct a UMAP representation. The data was preprocessed in R v4.1.0 with Seurat v4.1.0. Cells with unique feature counts below 200 or exceeding 6000 as well as cells with a mitochondrial count percentage exceeding 5% were excluded. Additionally, barcodes with UMI count below 200 or above 40000 were also removed. The data underwent log normalization with a scale factor of 10000, the top 2000 highly variable genes were identified by the vst method and scaled. Principal component analysis (PCA) was conducted on the scaled data and k-nearest neighbors were computed using the default Seurat settings and 40 principal components (PC) for FindNeigbors(). Subsequently, clusters identification by FindClusters() and UMAP visualization by RunUMAP() was conducted with a resolution of 0.8. After manual annotation of the clusters, differential expression analysis was carried out with FindAllMarkers() in default settings.

### Flow cytometry

#### GDF15-EEV titration: myeloid and lymphoid panels on spleen, MLN and gut

Spleen, the proximal three mesenteric lymph nodes (MLN), small intestine (SI) and colon were collected in RPMI medium on ice upon sacrifice. MLN and spleen were mashed with a plunger and then filtered using a 70µm filter. Red blood cells in spleen were eliminated using ACK lysis buffer (Westburg, Cat. 10-548E). The samples were centrifuged (350g, 5min, 4°C) and resuspended in complete RPMI medium for counting.

For gut tissues, lamina propria was isolated as follows: While keeping moist in 2% FCS HBSS, samples were cleaned, cut longitudinally and into 1cm segments. While in 10ml cold HBSS, the tissues were shaken vigorously to remove any debris, after which it was filtered through nylon. Next, 10ml of 2mM EDTA/HBSS was added, and the mixture was incubated with magnetic stirring at 37°C for 15min at 250rpm. After washing, this process was repeated, and the supernatants were removed each time using a new nylon mesh. Finally, the intestine was minced by cutting rapidly with scissors. 10ml enzyme cocktail, consisting of 0.425mg/ml Collagenase V (Sigma, Cat. C9263), 0.6mg/ml Collagenase VIII (Sigma, Cat. C2139-1G), 0.75mg/ml Collagenase D (Roche, Cat. 11088882001), and 50µg/ml DNase (Sigma, 1128492001), was added and incubated for 30min with magnetic stirring at 37°C, 250rpm. The resulting single cell suspension was filtered through a 100µm cell strainer, washed and resuspended using complete RPMI to count for staining.

Two million cells of each sample for each flow panel were stained as follows: Live/Dead staining (Fixable Viability Dye eFluor506, Thermofisher. Cat. 65-0866-14) for 10min at 4°C, then FcR (Miltenyi, Cat. 130-092-575) and monocyte (Biolegend, Cat. 2000135006) blocking for 10min at 4°C, followed by washing. Surface antibodies were added and incubated for 30min at 4°C in the dark. The surface staining mix for the lymphoid panel consists of Cell Staining Buffer (Biolegend, Cat. 420201), Brilliant Stain buffer (BD, 563794), CD127-AF700 (Thermofisher, Cat. 56-1271-80), CD45-BUV395 (BD, Cat. 564279), TCRgd-BV605 (Biolegend, Cat. 118129), CD3-BV711 (Biolegend, Cat. 100241), CD4-BV785 (Biolegend, Cat. 100452), NK1.1-PECy5 (Biolegend, Cat. 108716) I-A-APCeFluor780 (Thermofisher, 47-5321-80), CD19-APCeFluor780 (Life technologies, Cat. 47-0199-42), CD11b-APC.Cy7 (BD, Cat. 557657) and TCRb-BUV737 (BD, Cat. 564799). The myeloid panel consists of Cell Staining Buffer, Brilliant stain buffer, F4/80-biotin (Thermofisher, Cat. 13-4801-82), Ly6C-BV605 (Biolegend, Cat. 128035), CD64-BV711 (Biolegend, Cat. 139311), SIRPa-PE.Cy7 (Biolegend, Cat. 144008), CD206-APC (Biolegend, Cat. 141707), Ly6G-PerCPCy5.5 (BD, Cat. 560602), CD103-PE (Thermofisher, Cat. 12-1031-82), CD3-PECy5 (eBioscience, Cat. 15-0031-82), CD19-PECy5 (eBioscience, Cat. 15-0193-82), NK1.1-PECy5 (TONBO, Cat. 65-5941-U100), SiglecF-BUV395 (BD, Cat. 750280), CD11c-PE.eFluor610 (Thermofisher, Cat. 61-0114-82), I-A-BV785 (Biolegend, Cat. 197645), CD45-APCeFluor780 (eBioscience, Cat. 47-045180), XCR1-BV650 (Biolegend, Cat. 148220), CD11b-BUV737 (BD, Cat. 564443), CD86-FITC (BD, Cat. 553691). To the myeloid panel, Streptavidin-BV421 (BD, Cat. 563259) was added and incubated 15min at 4°C. All cells were washed twice with FACS buffer. Cells were fixed using the eBioscience TF fixative (Cat. 00-5523-00) according to the manufacturer’s instructions. For the lymphoid panel, cells were permeabilized and intracellular staining using Ki67-AF488 (BD, Cat. 561165), GATA3-BB700 (BD, Cat. 566643), RORgT-PE (Thermofisher, Cat. 12-6982-82), Tbet-PB (Biolegend, Cat. 644808) and FoxP3-AF647 (Biolegend, Cat. 126407) was performed by incubating 30min at room temperature. The cells were washed in permeabilization buffer, measured on a Fortessa (BD Biosciences) and analyzed using FlowJo.

#### Mouse adherent bone cell isolation, MALP staining and sorting

To isolate adherent bone marrow cells, we adapted previously published protocols^79,80^, so that antigens would be conserved for flow cytometry. In brief, all soft tissues were removed from both femurs and tibiae and the surface of the bones were scraped to remove the periosteum. The proximal and distal ends of the bones were cut off at the growth plate. Using a PBS-filled syringe, the non-adherent bone marrow was flushed until the bones were clear. The flushed bone marrow was kept for selected experiments where indicated. After rinsing, the bones were cut open longitudinally followed by transversally cutting into smaller pieces. The bones were digested in a sterile urine pot with PBS solution supplemented with 1mg/ml collagenase I (Thermofisher, Cat. 17018029), and 0.1mg/ml DNase I (Sigma, Cat. 11284932001) for 1hr at 37°C while magnetically stirring at 400rpm. The resulting cell suspension was filtered through a 100µm strainer and the red blood cells were eliminated using ACK lysis buffer. The cells were resolved in phenol red-free complete DMEM and counted for staining using trypan blue.

MALPs had not been detected by FACS to date. We first performed *in silico* identification of cell surface proteins by using DAVID (Database for Annotation, Visualization and Integrated Discovery; https://david.ncifcrf.gov/) to identify plasma membrane molecules from a list of MALP-specific genes identified by Zhong et al.^57^. We then searched the catalogs of antibody suppliers for FACS-validated antibodies against MALP-expressed cell surface genes. Candidate antibodies were screened in a panel to assess their expression on hematopoietic stem cells, immune cells, osteoblasts, osteocytes, and the remaining MALP-containing cells (as described below). Only Qa2 and CD106 (VCAM1) double-stained the remaining cells and were thus further explored as MALP markers for flow cytometry.

To validate Qa2+CD106+ cells as MALPs, we used FACS sorting and qPCR. Here, adhesive bone marrow cells were obtained using the above protocol and selected cell populations were sorted as follows: Live/Dead staining was performed using zombie NIR (Biolegend, Cat. 423106) for 30min at 4°C. After washing, the blocking mix, which included Fc block and monocyte block, was added, and incubated for 10min at 4°C. Then the surface antibodies were added and incubated for 30min at 4°C. The surface antibodies, which were diluted in cell-staining buffer containing BD brilliant buffer, included: CD90-BV510 (Biolegend, Cat. 140319), CD34-PE/Dazzle (Biolegend, Cat. 119329), CD117-APC (Biolegend, Cat. 105811), Sca1-BUV496 (BD, Cat. 750169), CD3-FITC (Biolegend, Cat. 155603), CD45-BV570 (Biolegend, Cat. 103135), CD106-BUV737 (Biolegend, Cat. 741726), Ter119-FITC (Biolegend, Cat. 116205), Qa2-biotin (BD, Cat. 558973), CD59a-PE (Biolegend, Cat. 143103), PDPN-PE-Cy7 (Thermofisher, Cat. 25-5381-82), CD45RB-FITC (Biolegend, Cat. 103305), CD11b-FITC (Invitrogen, Cat. 11-0112-41), CD31-BV711 (Biolegend, Cat. 102449) and Ly6G-FITC (Biolegend, Cat. 127605). Then, a secondary stain with streptavidin-BV421 was performed and incubated for 15min at 4°C. The cells were washed twice before being resuspended in preparation for sorting. The cell sorting was performed using a Symphony S6 (BD Biosciences). The cells were pelleted in 1.5ml tubes and the RNA was extracted for qPCR as described above.

As MALPs proved to be relatively rare cells (∼8000 in a total of 10 x 10^6^ adherent bone cells per mouse), long sorting durations were required for each sample. This made sorting of cells impractical for larger experiments. To speed up the process, we made the decision to enrich for MALPs using magnetic selection. This was achieved using the BioLegend MojoSort Mouse Hematopoietic Progenitor Cell Isolation Kit (Cat. 480004) and the Dynasort-15 magnetic holder. Briefly, after enzymatic digestion, the cells were washed with MojoSort buffer (Biolegend, Cat. 480017) and magnetic separation was performed according to the manufacturer’s instructions. The following changes were made to the protocol: 1) as the MojoSort antibodies and FITC antibodies (dump channel for lineage positive cells) of our panel bind to the same antigens, staining of these antibodies was performed simultaneously instead of during the later surface staining. 2) As the MojoSort antibodies are biotinylated we replaced the Qa2-bio antibody with Qa2-BV421 (BD, Cat. 743309). Using qPCR and FACS we ensured that the positively selected (lineage positive) cells did not contain Qa2+CD106+ MALPs. Using this protocol, we sorted ∼10^6^ negatively selected cells in which the MALP frequency was increased 10-fold over total adherent bone cells.

#### Human FACS

Biopsies processing: Human knee samples were obtained as pieces of bone removed during total knee replacement surgeries and smaller pieces of trabecular bone were further removed by 4mm punch biopsy tools (CLS-med). Biopsies were digested for 45-60min at 37°C in a digestion medium containing 5ml of HBSS (Sigma, Cat. H6638), 2.5mg of Collagenase P (Sigma, Cat. 11213857001), 500µg DNAse I (Sigma, Cat. 11284932001), and 500µg Liberase (Sigma, Cat. 5401020001). After digestion, cells were flushed from the bone tissue by syringe.

FACS protocol: To search for MALP-like cells in humans we utilized cryopreserved single-cell suspensions obtained from enzymatic digestion of biopsies taken from human knee samples received from knee-replacement surgeries of patients with osteoarthritis. We followed a strategy similar to that applied for MALPs identification by flow cytometry in mice. Live/dead staining was performed using Zombie NIR (Biolegend, Cat. 423106). We performed negative selection by firstly gating out cells of hematopoietic origin using CD71-BUV615 (BD, Cat. 743308) and CD45-PerCP (Biolegend, Cat. 304026), afterward excluding residual CD45^dim^ neutrophils by gating out CD11b^+^ cells (CD11b-SparkYG593, Biolegend, Cat. 101282), followed by gating out Podoplanin-positive (APC-Fire750, Biolegend, Cat. 337023) and ALP-positive cells (AlexaFluor647, BD, Cat. 561600) (osteocytes and osteoblasts respectively), cells expressing CD31 (BV605, Biolegend, Cat. 303121) and CD34 (BV785, Biolegend, Cat. 343625) (endothelial cells and hematopoietic stem cells) and finally cells expressing CD90 (BUV395, BD, Cat. 56804) (mesenchymal stromal cells) or CD117 (BV650, BD, Cat. 563859) (remaining progenitors). In the residual population, MALP-like cells were identified as the ADRB2^high^CD106^+^ population (using unconjugated ADRB2, Thermofisher, Cat. BS-0947R followed by rabbit IgG-FITC, Abcam, Cat. ab6717 and CD106-PE-Cy7, Biolegend, Cat. 305817). A 5-laser Cytek Aurora machine was used for the analysis.

To obtain sufficient numbers of rare MALP-like cells we pooled single cell suspension derived from two donors prior to sorting. Afterwards, we first performed MACS negative enrichment using BioLegend MojoSort Human anti-PE nanobeads (Cat. 480092) to sort out all lineage positive cells. Live/Dead staining was performed using Live/Dead Blue (Thermofisher, Cat. L23105). For the negative enrichment we used the same antibodies as for the staining but PE-labeled (Biolegend, Cat. 328109, 334105, 303105, 313203, 343605, 304058, 337004 and 305817; BD, Cat. 561433). After negative enrichment, we sorted MALP-like cells as ADRB2^high^CD106^+^ population. Other analyzed populations were sorted without previous enrichment using the same combination of antibody clones that were used for the staining but some conjugated with different fluorochromes to better fit the suit the sorter. We sorted CD45^+^CD71^-^ cells, osteoblasts (CD71^-^, CD45^-^, CD90^+^, ALP^+^), CD90^+^ALP^-^ subset (CD71^-^, CD45^-^, CD90^+^, ALP^-^), and endothelial cells (CD45^-^, CD71^-^, CD90^-^, ALP^-^, CD31^+^, CD34^+^). We used BD FACSAria™ for the sorting.

## QUANTIFICATION AND STATISTICAL ANALYSIS

Statistical analyses were conducted using GraphPad Prism 9 and GenStat 22. Residual plots were used to assess normal distribution of the residuals and to identify outliers. For each graph, the statistical analysis performed is discussed in the figure legends. All graphs show mean ± SEM and n is the number of individuals (i.e., patients or individual mice) per group.

Patients using metformin were excluded from the GDF15 serum analysis, as most of these patients were outliers, which has already been reported in the literature^81^. All patient data was corrected for age.

Simple linear regression with groups was used to test for parallelism: The first model to be fitted is a simple linear regression, ignoring the groups. Next the model is extended to include a different constant (or intercept) for each group, giving a set of parallel lines one for each group. Then, the final model has both a different constant and a different regression coefficient (or slope) for each group.

To avoid age-related differences in the case of in-house bred mice, multiple injection groups were pooled until we achieved the predetermined number of mice. The injection group was included as a factor in the statistical model specific for the read-out. For femur bending analysis, the width of the femur was included in this statistical model.

If only two groups were compared, we performed an unpaired t-test. To compare multiple independent groups, we used one-way ANOVA with post-hoc Tukey’s comparisons. Two-way ANOVA was performed to assess the significance between two factors (e.g., GENOTYPE and EEV). Measurements over time were analyzed as repeated measurements analysis using the Restricted Maximum Likelihood (REML) approach, accounting for the correlation structure between observations, and assessing the significance of e.g., GENOTYPE, EEV (where applicable), TIME, and its interaction. In this model, body weight data were adjusted for differences between baseline body weight. For statistics applied on RNA sequencing, please see the respective methods section.

## SUPPLEMENTAL TABLES TITLES AND LEGENDS

**Table S1. Accumulated analysis of variance tables for simple linear regressions with groups between serum GDF15 levels measured in DEXA-scanned RA patients and healthy controls, and various BMD parameters. Related to Figure 1**.

Serum GDF15 levels and bone mineral density (BMD; by DEXA scanning) were determined in RA (Rheumatoid Arthritis) patients and healthy controls. Simple linear regression with groups was used to test for parallelism. The statistical results are reported here. The factor “GROUP” represents the two patient groups: RA or healthy control. The reported p-values shown in Figure 1 are highlighted in bold and refer to the significance of the difference in regression coefficients between patient groups. d.f. = degrees of freedom; s.s. = sum of squares; m.s. = mean of sum of squares; v.r. = variance ratio; pr = probability.

**Table S2. Cohort 1 patient clinical characteristics, related to STAR methods**

Serum GDF15 was determined in 20 healthy controls, 20 RA patients and 111 SpA (Spondyloarthritis) patients. Patient clinical characteristics are shown. SJC = Swollen Joint Count; TJC = Tender Joint Count; ASDAS = Ankylosing Spondylitis Disease Activity Score; BASDAI = Bath Ankylosing Spondylitis Disease Activity Index.

**Table S3. Cohort 2 patient clinical characteristics, related to STAR methods**

Serum GDF15 was determined in 31 healthy controls and 110 PsA patients. Clinical characteristics are shown. DAPSA = Disease Activity index for PSoriatic Arthritis.

**Table S4. Cohort 3 patient clinical characteristics, related to STAR methods**

Serum GDF15 was determined in 29 PsO (psoriasis) patients. PASI = Psoriasis Area and Severity Index.

**Table S5. Cohort 4 patient clinical characteristics, related to STAR methods**

RA (Rheumatoid Arthritis) patients and healthy controls underwent DEXA scanning to determine bone mineral density (BMD). Serum GDF15 was determined. Patient clinical characteristics are shown. DAS28 = Disease Activity Score 28-joint count; SJC = Swollen Joint Count.

**Table S6, clinical score sheet for IL-23 EEV experiments, related to STAR methods**

Mice were injected with either IL-23 EEV or control EEV. Clinical psoriasis and arthritis were assessed and scored according to the table by two blinded investigators three times per week.

**Table S7, qPCR primer sequences, related to STAR methods**

qPCR primers were designed using the NCBI primer design tool and were tested for efficiency. The primer sequences used in experiments are shown.

## SUPPLEMENTAL FIGURES TITLE AND LEGENDS

**Figure S1. Increased GDF15 levels in arthritis associates with low bone density, related to Figure 1**

(A) Luminex was used to measure GDF15 in serum from HC (heathy controls, n=19) and SpA (Spondyloarthritis, n=111) patients. SpA patients were subdivided based on the presence (n=16) or absence (n=95) of skin or nail psoriasis. Data are represented by Tukey box and whiskers plots. A one-way ANOVA was used to test for differences between patient means, followed by a post-hoc Tukey test. Significances are indicated as: *p<0.05, ***p<0.001.

(B) Serum GDF15 was measured by ELISA and bone mineral density (BMD) assessed by DEXA scan in RA patients (n = 46) and HC (n = 54). Simple linear regression with groups was used to generate regression lines between serum GDF15 and femur neck BMD, inter-trochanter BMD, femur trochanter BMD and Ward’s triangle BMD, as indicated in the cartoon. The p-value refers to the significance of the difference in regression coefficients between patient groups HC and RA.

Each datapoint represents one patient. Serum GDF15 concentrations were adjusted for age. Significances are indicated as: *p<0.05, **p<0.01, ***p<0.001.

**Figure S2. GDF15 induces dose-dependent bone loss but no tissue inflammation, related to Figure 2**

(A) Schematic of the GDF15 Enhanced Episomal Vector (GDF15-EEV). The same vector lacking the GDF15 sequence was used as control, named “Control EEV”.

(B) Mice were injected with control EEV or increasing doses of GDF15-EEV. Serum was acquired at baseline and 1, 4 and 7 weeks after EEV administration. GDF15 levels determined using ELISA.

(C) Gating strategy showing live, single, CD45^+^ cells and cell frequencies for lymphoid cells in spleen.

(D) Gating strategy showing live, single, CD3^-^CD19^-^ cells and cell frequencies for myeloid cells in spleen.

(E) *Nfkb1* and *Stat3* counts determined by bulk RNAseq in liver, ankle, VAT (Visceral Adipose Tissue) and SI (Small Intestine), relative to control EEV.

(F) Inflammation-associated transcription factors in the liver determined by bulk RNAseq, relative to control EEV.

(G) Liver *Gdf15* counts determined by bulk RNAseq, relative to control EEV. A one-way ANOVA was used to test for differences in gene expression between the doses, followed by a post-hoc Tukey’s multiple comparison test.

(H) Bulk RNAseq on SI (small intestine, top) and ankle (including skin, muscle and tendon, bottom) of control EEV versus 1µg/5µg GDF15-EEV treated mice. Total number of differentially expressed genes (DEG) (log2 fold change >1.0 and <1.0, adjusted p value < 0.05).

(I) µCT images of tibia. Representative images shown, bone cortex in grey and trabeculae in blue. The indicated bone parameters were quantified. Bone density is the bone volume divided by total volume. Trabeculae volume is normalized to the total inside volume. A one-way ANOVA was used to test for differences between the four doses, followed by post-hoc test for trend. Data are represented as mean ± SEM, dots represent the individual mice. n.s. = not significant, *p<0.05, **p<0.01, ***p<0.001.

**Figure S3. GDF15 is not required for steady-state bone homeostasis, related to Results**

(A) CRISPR/Cas9 strategy to create GFRAL-KO mice. Using two gRNAs, we were able to delete 58 bp in exon 3 of the *Gfral* gene causing non-sense mediated decay. Right, gel electrophoresis of a GFRAL-KO mouse and a WT littermate.

(B) Bone marrow was isolated from GDF15-KO mice and WT littermates and cultured *in vitro* with RANKL and M-CSF to differentiate into osteoclasts over 5 days (n=6). Representative images of multinucleated osteoclasts are shown.

(C) Number of multinucleated cells counted and quantitative analysis of bone resorption, as measured by fluorescence released by cells derived from GDF15-KO mice and WT littermates is shown. NTC (negative control) are bone marrow cells stimulated with M-CSF, but not RANKL. An unpaired t-test was used to test for differences between GDF15-KO and WT.

(D and F) Representative µCT images of tibia of mice aged 14 weeks; GDF15-KO (orange) and GFRAL-KO (blue) on bottom. Within each sex, an unpaired t-test was used to test for differences between GDF15-KO and WT.

(E and G) GDF15-KO mice and WT littermates and GFRAL-KO mice and WT littermates body weight at 14-weeks of age. Within each sex, an unpaired t-test was used to test for differences between GDF15-KO and WT.

Results are represented as mean ± SEM, dots represent the individual mice.

**Figure S4. GDF15 is not required for non-inflammatory bone homeostasis, related to Results**

(A) GDF15-KO mice and WT littermates underwent ovariectomy (OVX) or sham surgery. A repeated measurements analysis was performed to assess overall changes over time in body weight between the four groups, represented by the interaction between genotype (KO or WT), surgery (OVX or Sham) and time. Body weights were adjusted for differences in baseline body weight.

(B) Representative µCT images of tibia shown. The indicated bone parameters were quantified. Femoral strength was assessed using a three-point bending assay. A two-way ANOVA was used to test for interactions between Genotype (KO or WT) and surgery (OVX or Sham).

(C) Serum GDF15 levels determined by ELISA tested for differences between sham or OVX surgery in WT mice.

(D) Body weight of aged (15-month-old) male GDF15-KO and WT littermate mice (n=13).

(E) Representative µCT images of tibia shown. The indicated bone parameters were quantified. (C, D, E) An unpaired t-test was used to test the difference between the groups.

Data are represented as mean ± SEM, dots represent the individual mice.

**Figure S5. The GDF15-GFRAL axis mediates IL-23 induced weight loss and trabecular bone loss, but not inflammation severity, related to Figure 5**

(A) C57Bl/6J mice were injected with either IL-23 EEV or control EEV. Serum IL-23 levels were determined using ELISA.

(B) A repeated measurements analysis was performed to assess overall changes over time in body weight between the two EEV groups IL-23 or control, represented by the interaction between EEV and time. Body weights were adjusted for differences in baseline body weight.

(C) A repeated measurements analysis was performed to assess overall changes over time in clinical scores between the two groups, represented by the interaction between EEV (IL-23 or control), and time.

(D) Representative µCT images of calcaneus shown. The indicated bone parameters were quantified. An unpaired t-test was used to test for difference between the treatment groups. Dots represent the individual mice.

(E) GDF15-KO and GFRAL-KO and their respective WT littermates were injected with IL-23 EEV or control EEV. Omental VAT (Visceral Adipose Tissue) weight upon sacrifice.

(F-G) Representative H&E-stained images (F) and statistical analysis (G) of histopathology. Green arrow indicates bone marrow edema in calcaneus. Orange arrow indicates infiltration in the synovial-enthesial complex.

(H) Quantitative analysis of calcaneus cortex determined by µCT. The indicated bone parameters were quantified using custom scripts.

(I-J) Representative µCT images of tibia shown, with the bone cortex in grey and trabeculae in blue. The indicated bone parameters were quantified.

(E, G, H, J) A two-way ANOVA was used to test for interactions between Genotype (KO or WT) and EEV (IL-23 or control). Results are represented as mean ± SEM. Significances of the interactions are indicated as: *p<0.05, **p<0.01, *** p<0.001.

**Figure S6. GDF15 activates MALPs to produce RANKL and M-CSF, related to Figure 6**

(A) Reconstructed single-cell dataset of digested bone from 3-month-old mice from Zhong et al.^35^, with expression of different adrenergic receptors. Cluster annotation shown in Figure 6A.

(B) Counts of osteoblast, immune cell, and HSC (hematopoietic stem cell) associated genes as determined in whole ankle RNAseq of mice treated with control EEV and mid– or high dose of GDF15-EEV. Each gene is normalized to control EEV.

(C) Relative gene expression in proximal tibia of mice treated with GDF15-EEV or control EEV determined by qPCR, normalized to reference genes *Gapdh* and *Pgk1* and relative to control EEV (n = 5-7). A one-way ANOVA was used to assess the effect of the EEV. Significance is indicated as: ****p<0.001.

(D) Gating strategy to identify MALPs in adherent bone cells. Live, single cells are shown. Selected cell populations shown in bold were sorted to determine gene expression.

(E) Gene expression in sorted adherent bone cells, normalized to reference gene *Gapdh* and relative to Lin+ cells.

(F) Adherent bone cells of mice treated with GDF15-EEV or control EEV analyzed using flow cytometry. Frequency (% of Lin-cells) of the indicated cell types is shown.

(G) *Kng2* expression relative to Lin^+^ in sorted cells treated with GDF15-EEV or control EEV.

(F, G) A two-way ANOVA was used, followed by a post-hoc Tukey’s multiple comparison test. Results are represented as mean ± SEM. Dots represent individual mice.

**Figure S7. MALP cells are present in human bones, related to Figure 6 and 7**

(A) UMAP of cell clusters defined in scRNAseq on enzymatically digested human vertebrae biopsies.

(B) Gating strategy to identify MALPs in enzymatically digested knee biopsies. Gated on live, single cells. Selected cell populations shown in bold sorted to determine gene expression.

(C) MFI (Mean Fluorescence Intensity) of selected surface markers (n=8). Each datapoint represents one patient.

(D) Projection of CD45^-^CD71^-^CD115^-^CD11b^-^ cells onto UMAP with cell populations indicated (top) and surface marker expression (bottom).

(E) MALPs and other major cell populations in the bone were sorted and gene expression was determined in two pooled human samples. Expression is normalized to reference gene *GAPDH*.

(F) Relative gene expression of the proximal tibia of mice treated with GDF15-EEV or control EEV and 6-OHDA or saline. The indicated genes are negative control genes, expressed by adherent bone cells, but not expressed by MALPs. Gene expression was normalized to reference genes *Gapdh* and *Pgk1* and relative to the “Control EEV – saline” group. Results are represented as mean ± SEM.

