## Supplementary figures and images for "GDF15 mediates inflammation-associated bone loss through a brain-bone axis"

### Supplemental figures

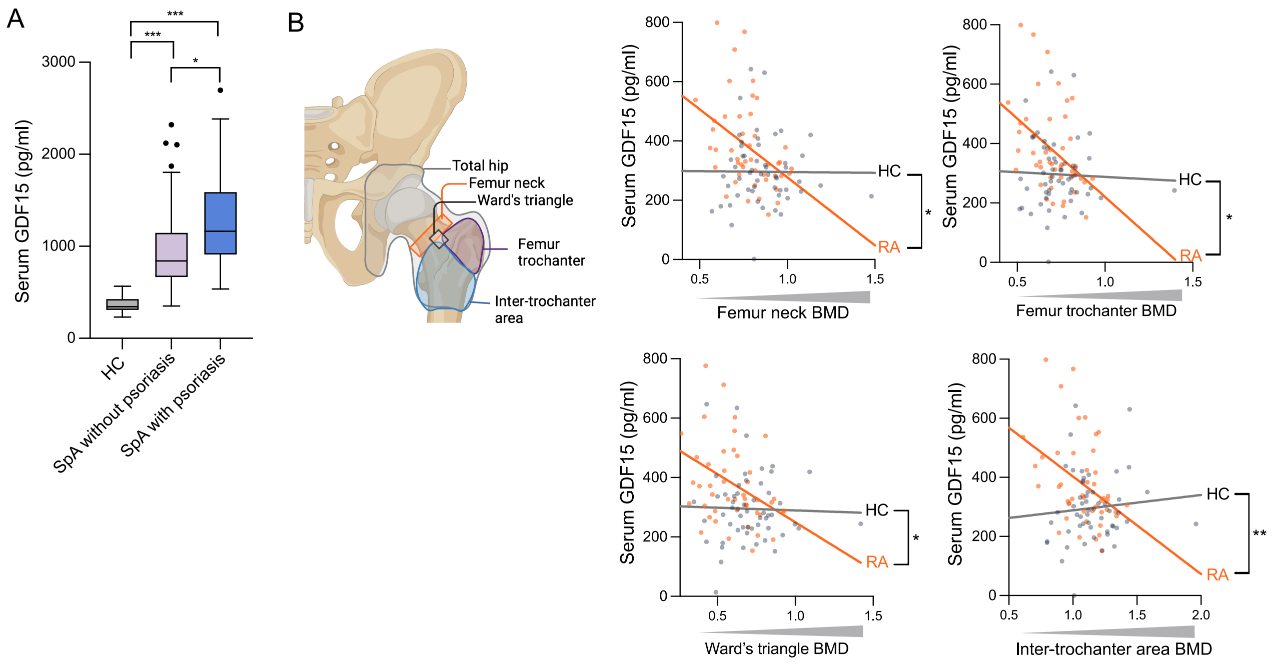

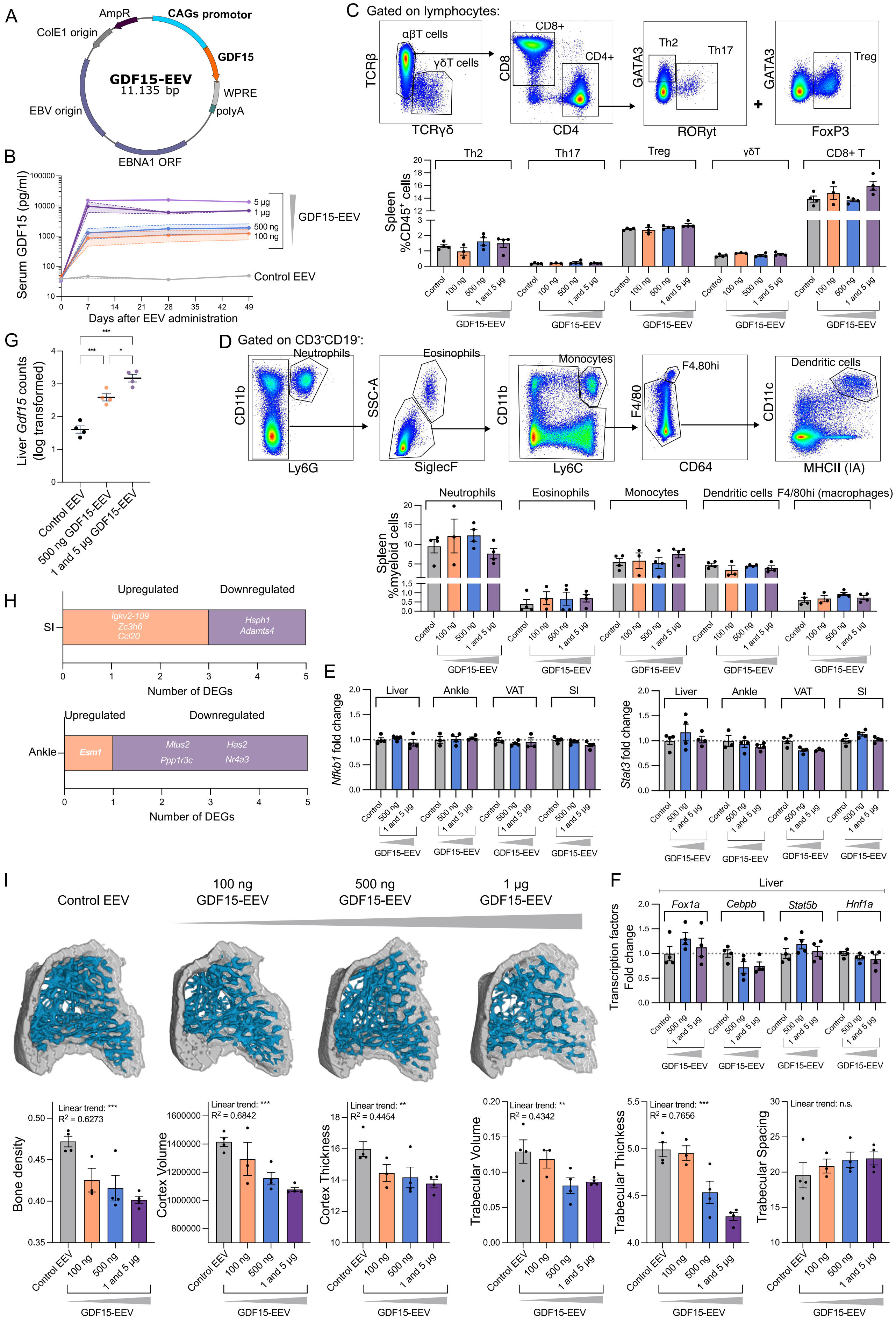

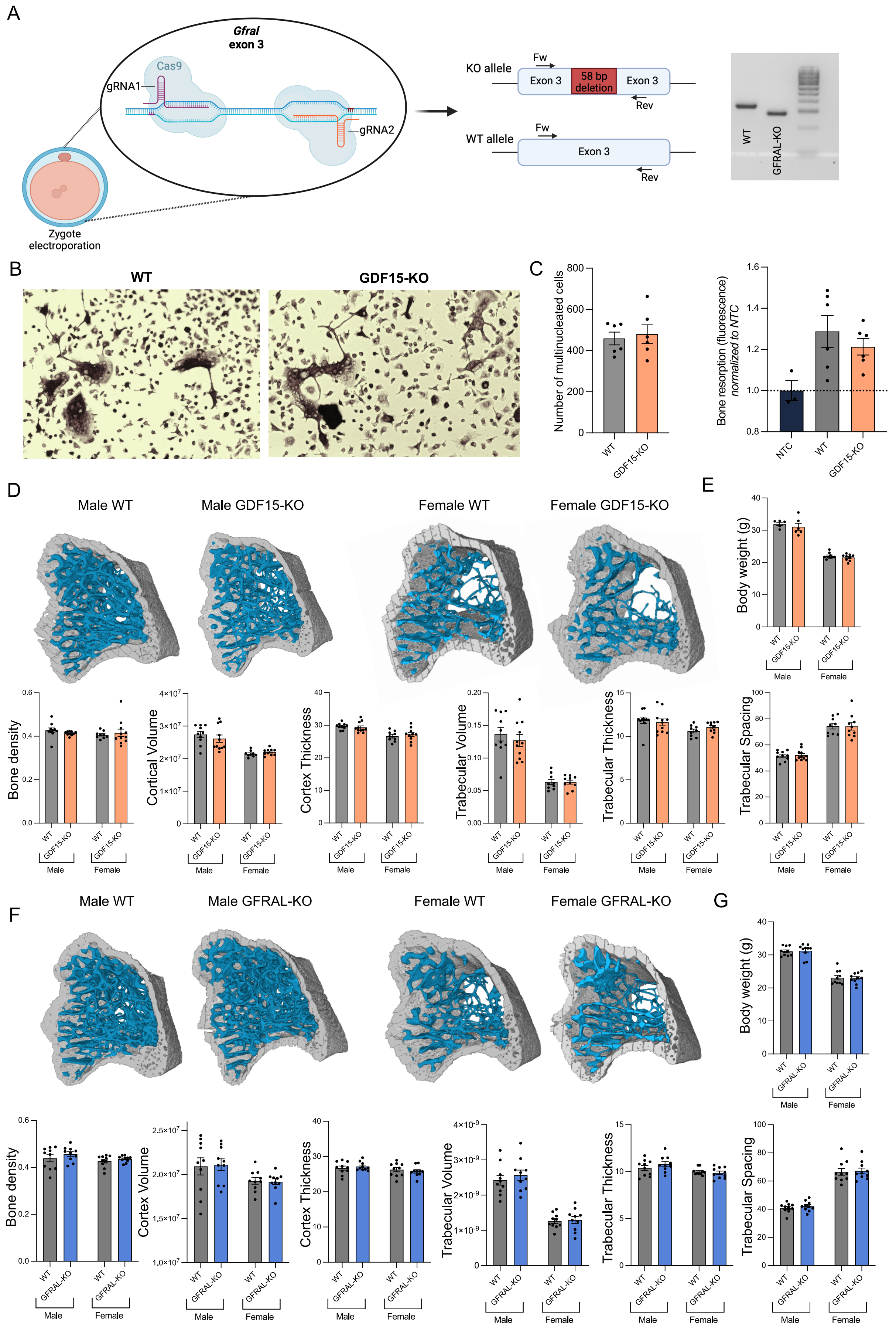

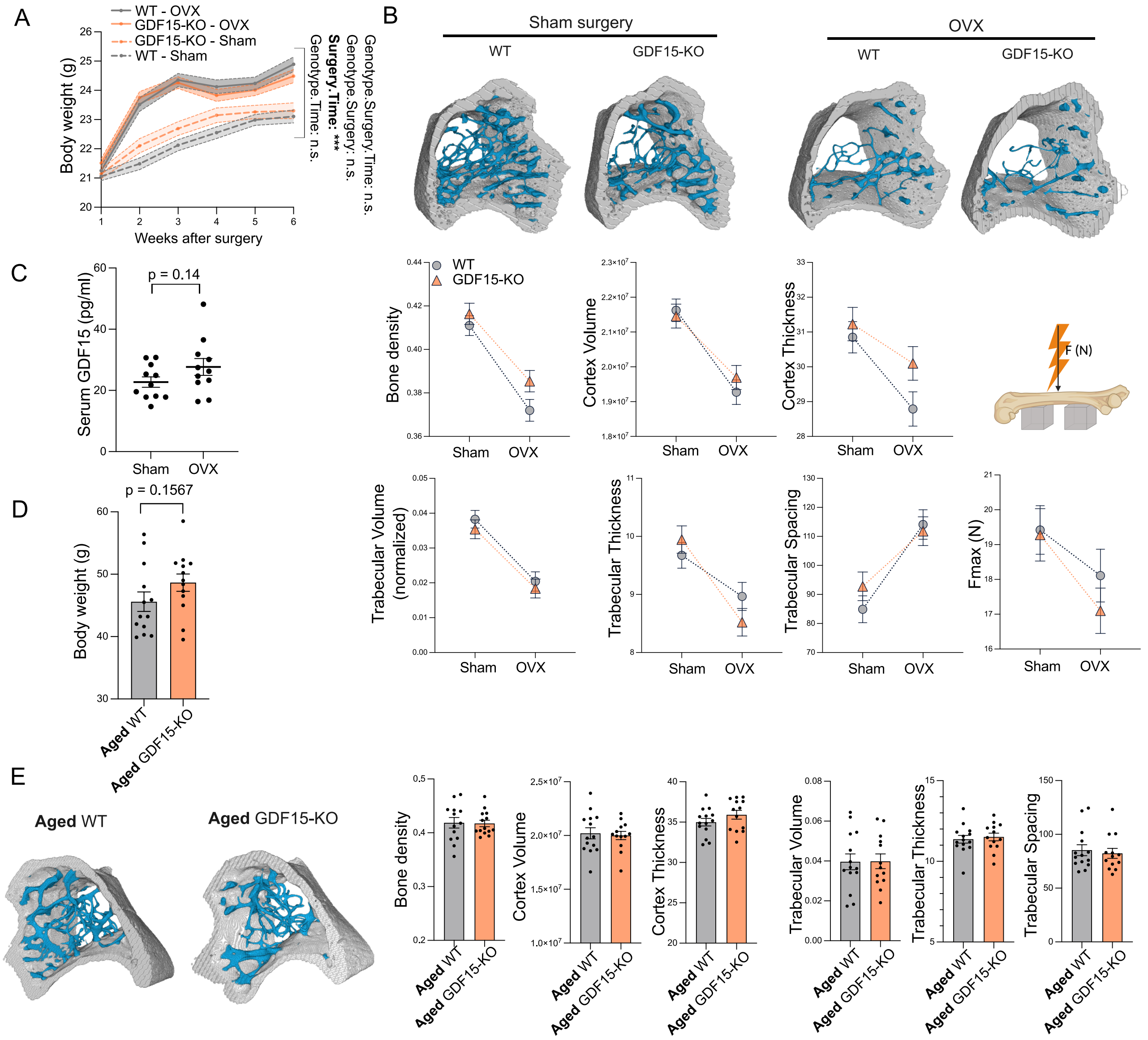

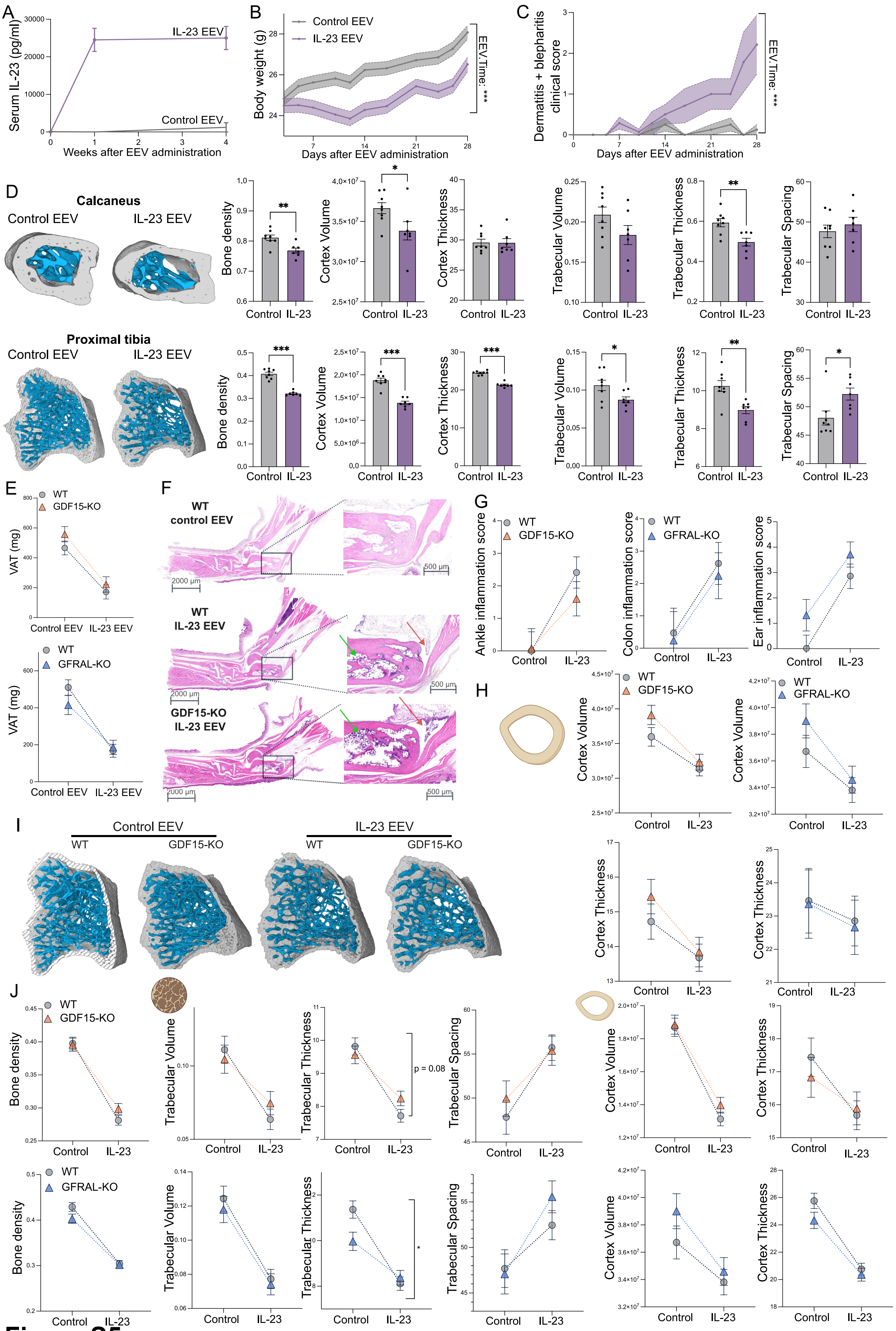

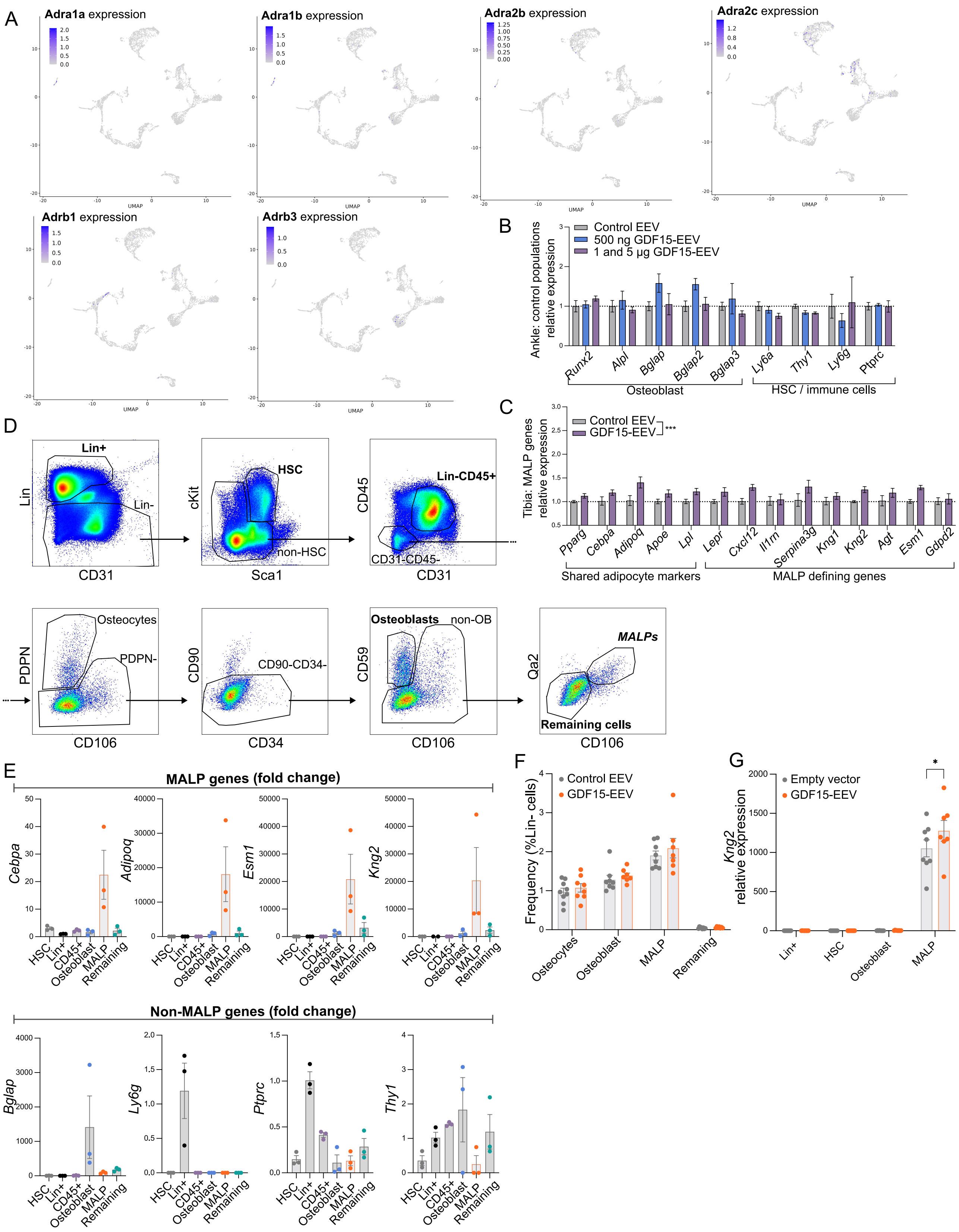

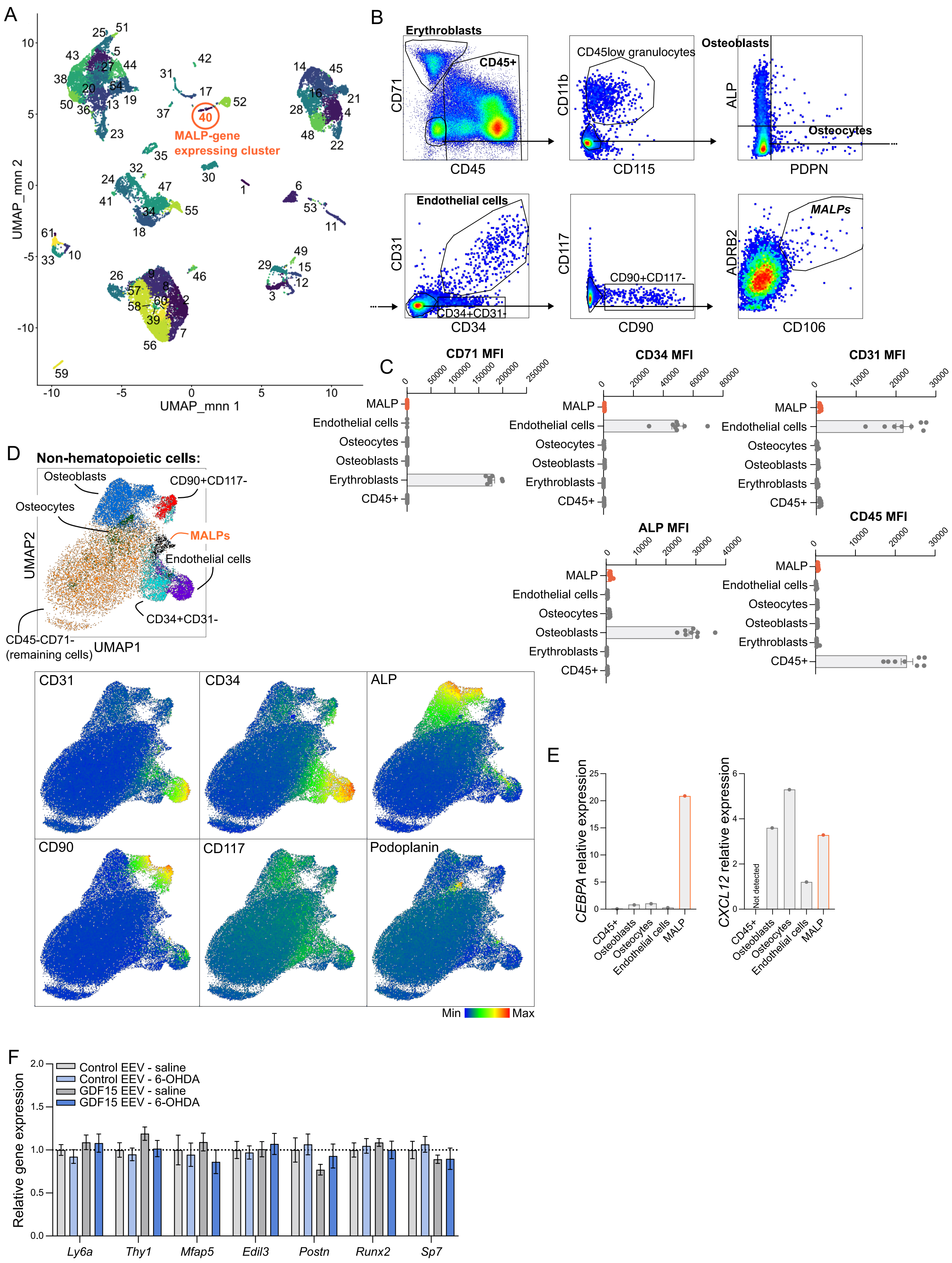
