## Supplemental tables for "GDF15 mediates inflammation-associated bone loss through a brain-bone axis"

**Table S1, Accumulated analysis of variance tables for simple linear regression with groups between serum GDF15 levels measured in DEXA-scanned RA patients and healthy controls, and various BMD parameters. Related to Figure 1.**

| Accumulated analysis of variance: **serum GDF15 vs total hip BMD** (HTOT_BMD) | | | | | |
|---|---|---|---|---|---|
| *Change* | *d.f.* | *s.s.* | *m.s.* | *v.r.* | *F pr.* |
| + age | 1 | 1356946 | 1356946 | 83.88 | <.001 |
| + group | 1 | 129795. | 129795. | 8.02 | 0.006 |
| + HTOT_BMD | 1 | 44593. | 44593. | 2.76 | 0.100 |
| **+ HTOT_BMD.GROUP** | 1 | 119750. | 119750. | 7.40 | **0.008** |
| Residual | 95 | 1536883. | 16178. | - | - |
| Total | 99 | 3187967. | 32202. | - | - |
| Accumulated analysis of variance: **serum GDF15 vs femur neck BMD** | | | | | |
| *Change* | *d.f.* | *s.s.* | *m.s.* | *v.r.* | *F pr.* |
| + age | 1 | 1356946 | 1356946 | 83.88 | <.001 |
| + group | 1 | 129795. | 129795. | 8.02 | 0.006 |
| + NECK_BMD | 1 | 60348. | 60348. | 3.72 | 0.057 |
| + **NECK_BMD.GROUP** | 1 | 99470. | 99470. | 6.13 | **0.015** |
| Residual | 95 | 1541408. | 16225. | - | - |
| Total | 99 | 3187967. | 32202. | - | - |
| Accumulated analysis of variance: **serum GDF15 vs inter-trochanter BMD** | | | | | |
| *Change* | *d.f.* | *s.s.* | *m.s.* | *v.r.* | *F pr.* |
| + age | 1 | 1356946 | 1356946 | 83.88 | <.001 |
| + group | 1 | 129795. | 129795. | 8.00 | 0.006 |
| + INTER_BMD | 1 | 35494 | 35494 | 2.19 | 0.142 |
| **+ INTER_BMD.GROUP** | 1 | 123942. | 123942. | 7.64 | **0.007** |
| Residual | 95 | 1541790. | 16229. | - | - |
| Total | 99 | 3187967. | 32202. | - | - |
| Accumulated analysis of variance: **serum GDF15 vs wards BMD** | | | | | |
| *Change* | *d.f.* | *s.s.* | *m.s.* | *v.r.* | *F pr.* |
| + age | 1 | 1356946 | 1356946 | 81.52 | <.001 |
| + group | 1 | 129795. | 129795. | 7.80 | 0.006 |
| + WARDS_BMD | 1 | 50062. | 50062. | 3.01 | 0.086 |
| **+ WARDS_BMD.GROUP** | 1 | 69817. | 69817. | 4.19 | **0.043** |
| Residual | 95 | 1581346. | 16646. | - | - |
| Total | 99 | 3187967. | 32202. | - | - |
| Accumulated analysis of variance: **serum GDF15 vs trochanter BMD** | | | | | |
| *Change* | *d.f.* | *s.s.* | *m.s.* | *v.r.* | *F pr.* |
| + age | 1 | 1356946 | 1356946 | 84.01 | <.001 |
| + group | 1 | 129795. | 129795. | 8.04 | 0.006 |
| + TROCH_BMD | 1 | 76038. | 76038. | 4.71 | 0.033 |
| **+ TROCH_BMD.GROUP** | 1 | 90718. | 90718. | 5.62 | **0.020** |
| Residual | 95 | 1534470 | 16152. | - | - |
| Total | 99 | 3187967. | 32202. | - | - |

**Table S2, cohort 1 patient clinical characteristics, related to STAR methods**

| | Healthy controls | RA | SpA |
|---|---|---|---|
| Total number (n) | 20 | 20 | 111 |
| Females (number) | 11 | 16 | 58 |
| Males (number) | 9 | 4 | 53 |
| Age (years) (mean □ SD) | 30.85 □ 6.60 | 56.2 □ 14.47 | 35.01 □ 10.54 |
| BMI (mean □ SD) | - | 28.88 □ 4.3 | 25.67 □ 4.59 |
| Anti-TNF treated (number) | - | 1 | 2 |
| CRP mg/l (mean □ SD) | - | 2.3 □ 7.16 | 1 □ 1.66 |
| SJC (mean □ SD) | - | 2.7 □ 3.757 | 0.9 □ 2.9 |
| TJC (mean □ SD) | - | 3.55 □ 4.86 | 1.86 □ 4.45 |
| Axial arthritis (number) | - | - | 71 |
| Peripheral arthritis (number) | - | - | 17 |
| Mixed axial/peripheral (number) | - | - | 23 |
| Psoriasis (number) | - | - | 16 |
| ASDAS (mean □ SD) | - | - | 4.35 □ 2.60 |
| BASDAI (mean □ SD) | - | - | 2.03 □ 0.92 |
| HLAB27+ (number) | - | - | 83 |

**Table S3, cohort 2 patient clinical characteristics, related to STAR methods**

| | Healthy controls | PsA |
|---|---|---|
| Total number (n) | 31 | 110 |
| Females (number) | 13 | 53 |
| Males (number) | 18 | 57 |
| Age (years) (mean □ SD) | 39.8 □ 8.42 | 49.37 □ 12.9 |
| BMI (mean □ SD) | - | 28.43 □ 5.53 |
| DAPSA (mean □ SD) | - | 13.44 □ 10.43 |
| CRP mg/l (mean □ SD) | - | 7.39 □ 6.02 |
| Biological treated (number) | - | 72 |

**Table S4, cohort 3 patient clinical characteristics, related to STAR methods**

| | **PsO** |
|---|---|
| Total number (n) | 29 |
| Females (number) | 12 |
| Males (number) | 17 |
| Age (years) (mean □ SD) | 45.03 □ 16.65 |
| Obese patients (number) | 8 |
| PASI (mean □ SD) | 9.24 □ 6.88 |
| Biological experienced (number) | 9 |
| Patients with a history of joint complaints (number) | 3 |
| Patients with joint complaints at time of blood draw (number) | 0 |

**Table S5, cohort 4 patient clinical characteristics, related to STAR methods**

| | **Healthy controls** | **RA** |
|---|---|---|
| Total number (n) | 54 | 46 |
| Females (number) | 29 | 33 |
| Males (number) | 34 | 20 |
| Age (years) (mean □ SD) | 46.62 □ 13.24 | 55.88 □ 12.20 |
| BMI (mean □ SD) | 24.55 □ 3.82 | 25.94 □ 4.79 |
| DAS28 (mean □ SD) | - | 5.53 □ 8.51 |
| CRP mg/ml (mean □ SD) | - | 14.38 □ 17.23 |
| SJC (mean □ SD) | - | 4.96 □ 4.82 |
| Biological-treated (number) | - | 6 |

**Table S6, clinical score sheet for IL-23 EEV experiments, related to STAR methods**

| What | How | Score |
|---|---|---|
| **Arthritis** | Toes: 0 = no swelling; 0.5 = 1 toe; 1 = 2 or more toes<br>PLUS Foot/hand: 0 = no swelling; 1 = swelling<br>PLUS Ankle/wrist: 0 = no swelling; 1 = swelling | 0-3 |
| **Blepharitis (eye)** | Can be seen in rim around the eye: hair loss, scaling, thickening.<br>0 = nothing; 0.5 = 1 eye mild; 1 = mild in both eyes; 1.5 = 1 eye severe + 1 eye mild; 2 = severe both eyes | 0-2 |
| **Dermatitis snout** | Hair loss, scaling, reddening.<br>0 = nothing; 0.5 = suspected thickening; 1 = thickening and scaling; 1.5 = clear hair loss towards eyes; 2 = clear hair loss (almost) reaching the eyes | 0-2 |
| **Dermatitis ear** | Scaling, thickening, reddening.<br>0 = nothing; 0.5 = thickening only 1 ear; 1 = thickening both ears with minor scaling; 1.5 = clear thickening and scaling both ears; 2 = clear thickening and severe scaling both ears. | 0-2 |
| **Dermatitis paws** | Scales on paws (to be seen between week 1 and 2).<br>Overall: score = number of paws that show dermatitis. 1 = 1 paw with clear dermatitis or 2 paws with very mild dermatitis | 0-4 |
| **Dermatitis tail** | White lines and scaling on tail.<br>0 = nothing; 0.5 = suspected very mild lines and/or scaling; 1 = moderate amount of lines and mild scaling; 1.5 = lines over entire length of tail and moderate amount of scales; 2 = clear lines and severe scaling over entire length of tail | 0-2 |

**Table S7, qPCR primer sequences, related to STAR methods**

| Gene | Forward primer OR Taqman probe ID | Reverse primer |
|---|---|---|
| *Gfral* | TTCCTGGCTGTTACGTTAAGC | GCCATTTGCATCAATCAAGCA |
| *Pparg* | TCACAATGCCATCAGGTTTGG | CTGGGTTCAGCTGGTCGATA |
| *Cebpa* | TTCGGGTCGCTGGATCTCTA | CCCTCATCTTAGACGCACCG |
| *Adipoq* | ATCTGGAGGTGGGAGACCAA | GGGCTATGGGTAGTTGCAGT |
| *Apoe* | TCGAGTGGCAAAGCAACCAA | TCAGTGCCGTCAGTTCTTGT |
| *Lpl* | TGCCCTAAGGACCCCTGAAG | ACATTCCCGTTACCGTCCATC |
| *Lepr* | ATGGTCACCCAGCACAATCC | GACGCAGTTTTTGGGCTCAG |
| *Cxcl12* | AAACTGTGCCCTTCAGATTG | GCCTCTTGTTTAAAGCTTTCTC |
| *Il1rn* | TAGTGTGTTCTTGGGCATCCA | TCAGTGATGTTAACTTCCTCCAG |
| *Serpina3g* | GAACCCCTTTGACCCGAATGA | GGGAGTTGTCAGGTAACCAGTTT |
| *Kng1* | TTGCACCGAGTGATCGAGG | TCAGCGTCCTTGAAGTCACA |
| *Kng2* | CACTGGAGAATGCACAGCAAC | ACAACTCGCACAGTGGTACTC |
| *Agt* | ACCCAGTTCTTGCCACTGAG | AAACCTCTCATCGTTCCTTGG |
| *Esm1* | CCGTACAGTCTCAGGCATGG | CCATCCCGAAGGTGCCATAG |
| *Gdpd2* | CATTGGGGACACTGGATGGA | CCCAGGAGCAATAGCAAGGAT |
| *Tnfsf11* | TGATGGAAGGCTCATGGTTGG | GACTTTATGGGAACCCGATGG |
| *Csf1* | AGACCAGGAACAGCTGGATGA | TCTCGGTGGCGTTAGCATTG |
| *Adrb2* | TCGTGCACGTTATCAGGGAC | GTTGACGTAGCCCAACCAGT |
| *Bglap* | CCCTGAGTCTGACAAAGCCT | ACTGAAGCTCCAAGGTAGCG |
| *Ly6g* | GGGAGGGGCTGAGAGAAAGTA | AGGGCTGCACAGATAAAACTTCC |
| *Ptprc* | TCCTCGTCCACTGCAGAGAT | CCGTGCTTTGCGTAGAGACT |
| *Thy1* | ACCAAGGATGAGGGCGACTA | ACACTTGACCAGCTTGTCTCTA |
| *Gapdh* | ACCCCAGCAAGGACACTGAGCAAG | TGGGGGTCTGGGATGGAAATTGTG |
| *Pgk1* | CTCCGCTTTCATGTAGAGGAAG | GACATCTCCTAGTTTGGACAGTG |
| *CEBPA* | AAGCACGATCAGTCCATCCC | AGACGCGCACATTCACATTG |
| *ADIPOQ* | GTATCCCCAAGCCACACCAT | TACACTTGCTGGGGCATCTG |
| *ESM1* | TGTGACAGCAGTGAGTGCAA | TGAGACTGTGCGGTAGCAAG |
| *PTPRC* | CAACAGTGGAGAAAGGACGC | GGCAAAGCCAAATGCCAAGA |
| *GAPDH* | Taqman: Hs02786624_g1 | - |
| *SERPINA3* | Taqman: Hs00153674_m1 | - |
| *CXCL12* | Taqman: Hs03676656_mH | - |
